# Polymorphic Structures of an Artificial Nucleocapsid Evolved from a Circularly Permuted Bacterial Protein Cage

**DOI:** 10.64898/2026.10.23.756749

**Authors:** Hualin Li, Łukasz Koziej, Osamu Kikko, Kazuki Ishihara, Takeru Kagawa, Kenya Tajima, Hideki Shigematsu, Christoph Gerle, Tomoaki Matsuura, Tomoyuki Numata, Yusuke Azuma, Naohiro Terasaka

## Abstract

Molecular self-assembly provides a route to nanoscale structures with defined sizes and shapes. Protein cages assembled from repeated subunits form compartments that can package molecular cargoes. Although amino acid substitutions can alter intersubunit interactions and cage architecture, how protein backbone connectivity relates to the cage architectures accessible during laboratory evolution remains unclear. Here we evolved an artificial nucleocapsid from a circularly permuted variant of *Aquifex aeolicus* lumazine synthase (AaLS). Laboratory evolution increased the yield of purified nucleocapsids relative to the progenitor, and the evolved variant, NC119-3, packaged its encoding mRNA. Cryo-electron microscopy revealed a heterogeneous population of cages, within which we esolved 180- and 210-subunit cage families composed of 36 and 42 pentamers, respectively. Within each resolved family, particles with the same subunit stoichiometry differed in the arrangement and orientation of their pentameric building blocks. A 2.90 Å reconstruction of a tetrahedral 180-subunit cage further revealed that sequence-identical subunits adopt distinct conformations depending on their positions within the assembly. These pentamer-based architectures differ from the domain-swapped *T* = 4 cage previously reported for nucleocapsids evolved from an AaLS variant with a different circular permutation site. Together with previous work, these findings show that RNA-packaging nucleocapsids evolved from scaffolds with different backbone connectivity can adopt distinct cage architectures.

**Contents Graphic:** 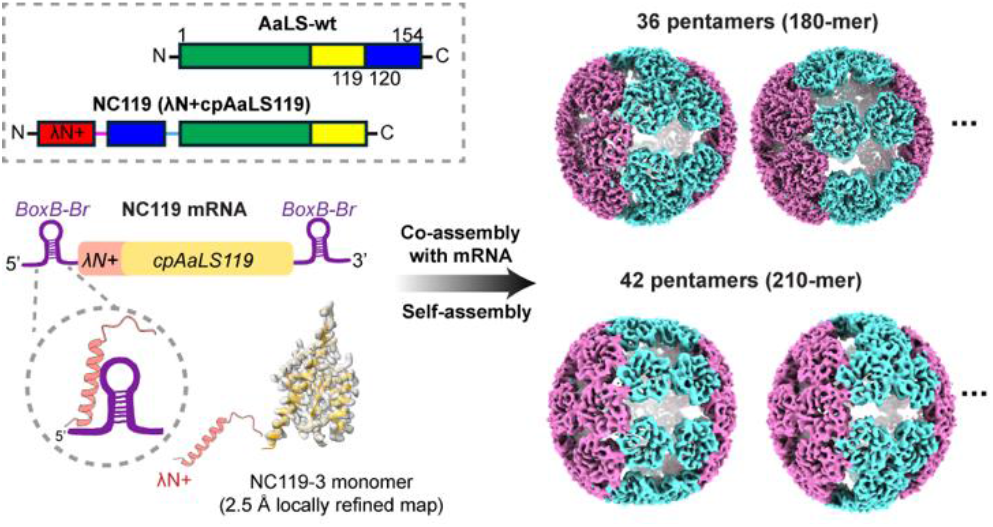

## Introduction

Protein cages consist of repeated protein subunits that self-assemble into closed compartments with defined sizes and internal volumes. ^1, 2^ Identifying how subunit conformation and intersubunit nteractions determine cage architecture can help explain the structural diversity of natural protein cages and guide the engineering of artificial nanocompartments. Amino acid substitutions and directed evolution can alter these conformations and interactions, giving rise to different cage architectures. ^3–5^ Altering polypeptide backbone connectivity provides another way to modify protein structure and assembly. Circular permutation, in which the original termini are linked and new termini are introduced elsewhere in the sequence, has altered the oligomeric organization of a bacterial shell protein and enabled peptide display and protein encapsulation in virus-like particles. ^6, 7^ Such changes in backbone connectivity can also provide different starting points for directed evolution. For example, directed evolution improved the activity of a circularly permuted β-lactamase. ^8^ Recently, directed evolution and topology reversion of a protein catenane showed that protein topology can modulate the effects of sequence variation on enzyme activity and stability. ^9^ How polypeptide backbone connectivity influences the structural repertoires reached during functional evolution of protein cages remains less clear.

*Aquifex aeolicus* lumazine synthase, AaLS, provides a useful system for studying this question. Native AaLS forms a 60-subunit icosahedral cage from 12 pentameric building blocks. ^10^ AaLS has been extensively engineered by amino acid substitution and laboratory evolution. ^5, 11^ These modifications have produced expanded cages with architectures distinct from the native 60-subunit structure, demonstrating structural plasticity of this protein cage. ^5^ Backbone connectivity has also been altered by circular permutation, in which the native termini are linked and new termini are introduced at another position in the sequence. ^12^

A circularly permuted AaLS variant with new termini between residues 84 and 85, termed cpAaLS84, was previously used as the starting point for evolving artificial nucleocapsids that package their own mRNA. ^13^ This variant forms a uniform 60-subunit cage. To introduce RNA-packaging ability, the cationic λN+ peptide was fused to its new *N*-terminus to generate NC-1 (λN+cpAaLS84) (Figure 1A, B). The peptide recruits the corresponding mRNA bearing engineered BoxB-Br packaging signals. NC-1 and the subsequently evolved NC-2 variant formed heterogeneous particles. Further evolution yielded NC-3, which adopted a 240-subunit icosahedral architecture with a triangulation number of 4 (*T* = 4), involving non-native domain-swapping between constituent capsomers. ^14, 15^ NC-4 retained the same architecture as NC-3 and showed improved and more selective packaging of full-length mRNA, accompanied by the evolution of an RNA packaging-signal cassette. ^15^ These studies demonstrated that extensive structural reorganization can occur during the evolution of an artificial nucleocapsid. They also raised the question of how the backbone connectivity of the starting protein influences the assembly states accessible during functional evolution.

**Figure 1.**
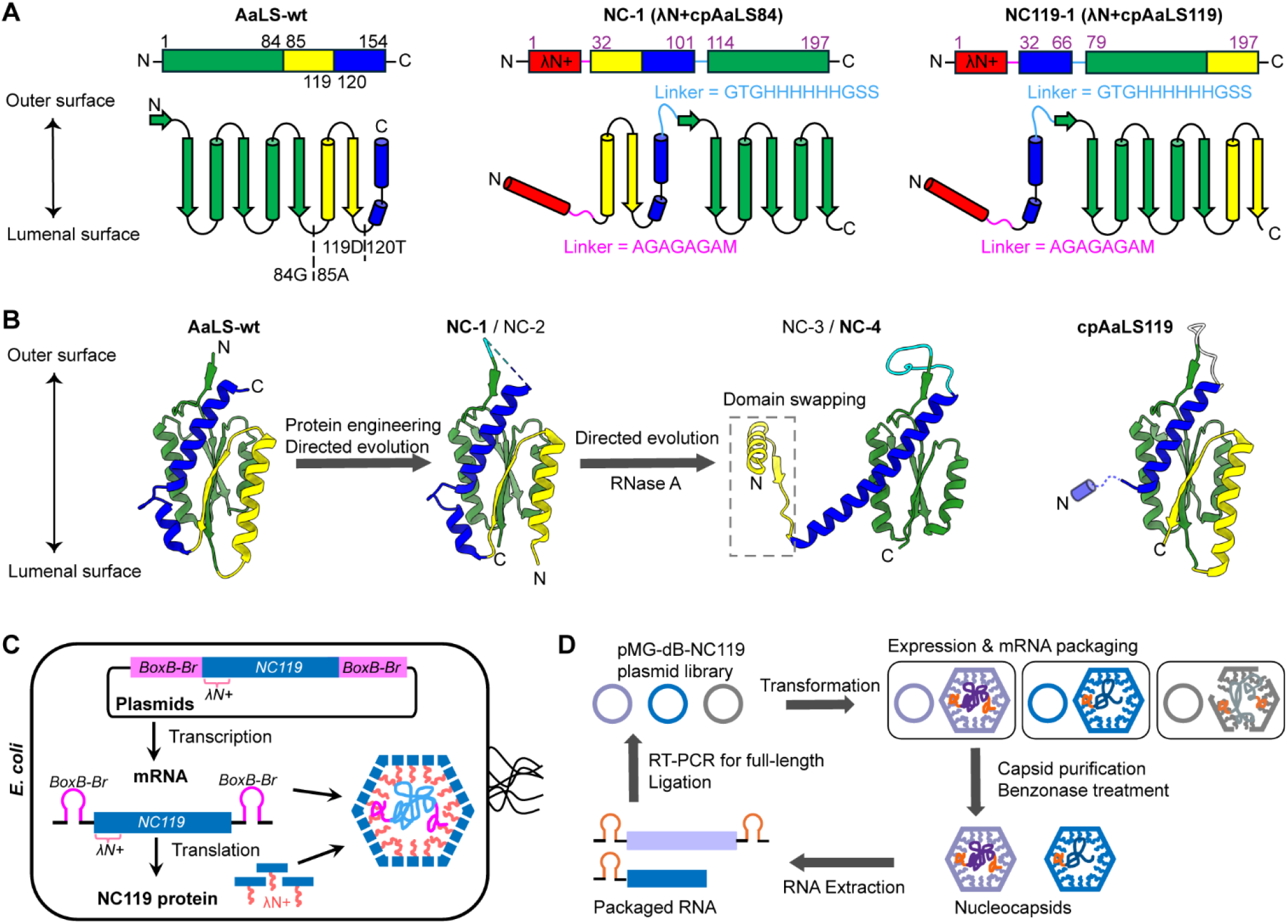
Topological design and directed evolution of the cpAaLS119 nucleocapsid lineage. **A)** Linear and topology diagrams of wild-type AaLS (AaLS-wt), the λN+ fused cpAaLS84 construct NC-1), and the λN+-fused cpAaLS119 construct (NC119-1). **(B)** Structural comparison of monomers rom AaLS-wt (PDB 5MPP), the previously reported NC-1 (PDB 7A4G) and domain-swapped NC-4 monomer (PDB 7A4J), and cpAaLS119 (PDB 9G3N). The evolution step from NC-2 to NC-3 included selection with RNase A. The dashed box identifies the region involved in domain swapping. The blue cylindrical cartoon in cpAaLS119 indicates a region unresolved in the reported structure (residues 120– 132 in AaLS-wt). The λN+ peptides are not shown in the NC-1 and NC-4 structures because they were not resolved in the corresponding structural models. **(C)** Schematic illustration of expression and RNA packaging in *E. coli*. BoxB-Br elements flank the NC119 coding region, and lumen-facing λN+ peptides recruit the mRNA during cage assembly. **(D)** Schematic illustration of the directed evolution workflow. *E. coli* cells are transformed with the pMG-dB-NC119 plasmid to express nucleocapsids. Purified nucleocapsids are treated with Benzonase to degrade unprotected RNA. Packaged RNA is extracted, and full-length cDNA is recovered by RT-PCR and ligated into the expression vector to generate the next-generation plasmid library.

We therefore examined cpAaLS119, which contains new termini between residues 119 and 120. This unction lies within the region corresponding to residues 85–154 of wild-type AaLS that participates in domain swapping in the evolved cpAaLS84-derived nucleocapsids NC-3 and NC-4 (Figure 1A, B). ^15^ introducing new termini within this region changes its backbone connectivity and may alter its ability to adopt the same domain-swapped arrangement. Previous studies have also shown that cpAaLS119 is structurally polymorphic, forming spherical cages and nanotubes depending on the linker connecting the native termini and on the solution conditions. ^12, 16^

Here, we report the laboratory evolution and structural characterization of a cpAaLS119-derived artificial nucleocapsid lineage, designated NC119. The evolved variants NC119-2 and NC119-3 both yielded more assembled nucleocapsids after purification than the initial NC119-1 construct and packaged their own encoding mRNAs. Cryo-electron microscopy of NC119-3 revealed a heterogeneous population containing 180- and 210-subunit cage families as well as larger cages whose structures remain unresolved. The resolved 180- and 210-subunit cages were built exclusively from pentamers. Within each of these two cage families, particles with the same subunit stoichiometry differed in the geometric arrangement of their pentameric building blocks. High-resolution analysis of a tetrahedral 180-subunit cage further revealed that sequence-identical subunits adopt distinct conformations depending on their positions within the assembly. These findings show that functional evolution from cpAaLS119 can yield RNA-packaging nucleocapsids, with NC119-3 adopting multiple pentamer-based architectures distinct from domain-swapped *T = 4* architecture previously characterized in the cpAaLS84 lineage.

## Results and Discussion

### Laboratory evolution restored productive assembly of the NC119 nucleocapsid

We first fused λN+peptide to the *N*-terminus of cpAaLS119 to generate NC119-1 (Figure 1A, B). BoxB-Br cassettes were introduced into both the 5′ and 3′ untranslated regions of the NC119-1 mRNA to promote λN+ peptide-mediated mRNA recruitment during intracellular assembly (Figure 1C). NC119-1 was expressed in *Escherichia coli* and purified by immobilized metal affinity chromatography (IMAC) and size exclusion chromatography (SEC). Cage-like particles approximately 30 nm in diameter were observed by negative-stain transmission electron microscopy (nsTEM), but nucleocapsid recovery after purification was low (Figure S1A, B). To improve productive nucleocapsid assembly, we generated a randomly mutagenized NC119-1 library by error-prone PCR and designated it NC119-1-Lib1. Following expression in *E. coli* and IMAC purification, the nucleocapsid library was treated with endonuclease Benzonase) to degrade unprotected RNA (Figure 1D and S1C, D). The remaining RNA was extracted and reverse transcribed. The resulting cDNA was amplified by PCR and ligated back into the expression vector to generate the next-cycle library, NC119-1-Lib2. Upon expression and purification, NC119-1-Lib2 showed a higher A260/A280 ratio in the cage-sized SEC peak than NC119-1-Lib1 (Figure S1C, E), and SDS-PAGE showed a predominant band corresponding to the nucleocapsid protein in the IMAC eluate and cage-sized SEC fraction (Figure S1F). Spherical particles were observed in this fraction by nsTEM (Figure S1E).

After two additional cycles to generate NC119-1-Lib4, 10 clones were arbitrarily chosen and sequenced Figure S2 and Table S1). One clone retained the parental NC119-1 sequence, whereas the remaining nine clones represented seven unique RNA sequences. Individual variants were examined by nsTEM, revealing cage-like particles in several variants (Figure S3). NC119-1-#02 formed particles spanning a broad range of sizes. We selected this variant for the next round of evolution to explore whether the cpAaLS119 scaffold could support larger and more structurally diverse nucleocapsids and designated t NC119-2. NC119-2 carried six amino acid substitutions relative to NC119-1, R7C, A51V, H73Q, H74L, L183S, and T195S (Figure 2A).

**Figure 2.**
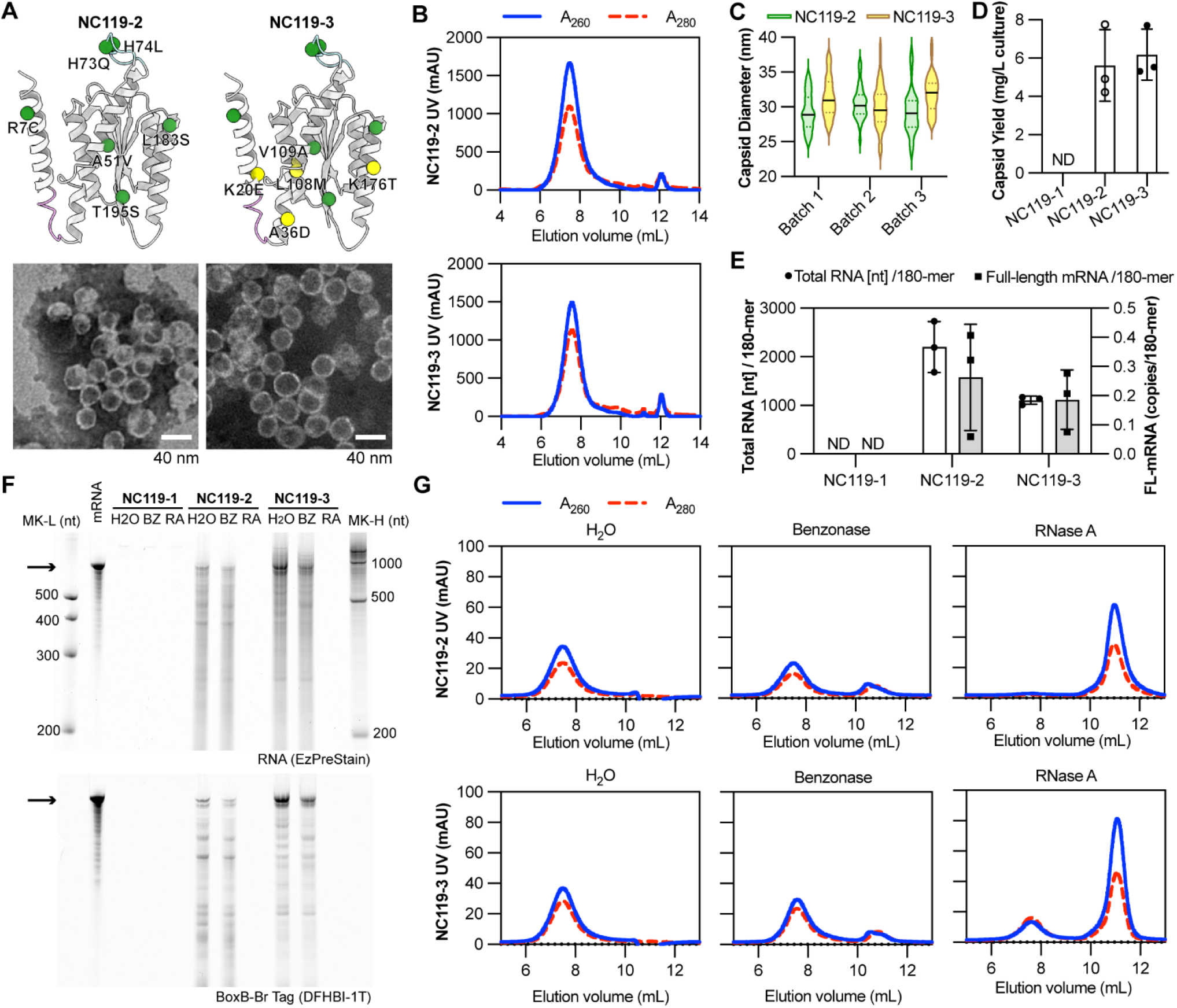
Biochemical and morphological characterization of NC119 variants. **A)** AlphaFold 3-predicted structures showing mutations acquired during evolution. Mutations in NC119-2 relative to NC119-1 and additional mutations in NC119-3 relative to NC119-2 are shown as green and yellow spheres, respectively. Representative negative-stain TEM images are shown below (scale bars, 40 nm). **(B)** Size exclusion chromatograms of IMAC-purified NC119-2 and NC119-3. Solid blue and dashed red lines indicate absorbance at 260 and 280 nm, respectively. **(C)** Particle diameters measured by nsTEM from three independent preparations (Batch 1–3), plotted separately. Green and yellow violin plots denote NC119-2 and NC119-3, respectively; dotted lines indicate the quartiles, and solid black lines indicate the median. **(D)** Purified nucleocapsid yields, expressed as the total mass of recovered protein–RNA complexes per liter of culture. ND, not detected. **(E)** Total RNA content (left axis, circles and open bars) and full-length mRNA copy numbers (right axis, squares and gray bars) packaged, normalized per 180-subunit equivalent. In (D) and (E), bars show mean ± SD from three independent preparations per variant. NC119-2 and NC119-3 were compared using two-tailed unpaired Welch’s *t*-tests: yield, *p* = 0.6967; total RNA, *p* = 0.0657; FL-mRNA, *p* = 0.5683. **(F)** Denaturing urea-PAGE of extracted RNA from nucleocapsids after water (H_2_O), Benzonase (BZ), or RNase A (RA) treatment. EzPreStain staining shows total RNAs (top), and DFHBI-1T staining shows BoxB-Br-containing RNA (bottom). MK-L and MK-H denote low- and high-range RNA size markers, respectively. Arrows indicate the expected positions of full-length NC119 mRNA. For the H₂O- and Benzonase-treated samples, 1,000 ng of extracted RNA was loaded per lane when available; otherwise, the maximum available amount was loaded. **(G)** Size exclusion chromatograms of NC119-2 (top) and NC119-3 (bottom) after treatment with water, Benzonase, or RNase A.

### A second round of evolution yielded NC119-3 with a broad particle-size distribution

NC119-2 was diversified by error-prone PCR to produce the next generation library, NC119-2-Lib1. Three cycles of Benzonase-based selection were performed to enrich variants capable of assembling nucleocapsids that protect encapsulated RNA from nuclease digestion. Following selection, twenty-five individual clones were isolated and sequenced (Table S2). Four clones retained the parental NC119-2 sequence, whereas two additional clones contained only synonymous substitutions and therefore encoded the parental NC119-2 protein sequence. The remaining 19 clones contained one or more missense mutations. These 19 variants were expressed individually in *E. coli*, purified by IMAC, and visualized by nsTEM to assess particle assembly and morphological variation (Figure S4). Five assembled variants were re-examined by nsTEM and further purified by SEC (Figure S5). Among these, NC119-2-#05 showed distinct particles in two independent expression experiments (Figure S4 and S5), including larger particles approaching 40 nm in diameter. It also exhibited a prominent cage-sized SEC peak and a minor peak corresponding to disassembled cage fragments showing highly efficient nucleocapsid assembly (Figure S5). Based on these observations, NC119-2-#05 was selected for subsequent biochemical and structural characterization and designated NC119-3. NC119-3 contained ive additional amino acid substitutions relative to NC119-2: K20E, A36D, L108M, V109A, and K176T Figure 2A and S6).

Both NC119-2 and NC119-3 produced distinct cage-sized SEC peaks, and nsTEM revealed intact but morphologically heterogeneous particles (Figure 2A, B). Particle diameters were determined from nsTEM images by averaging two perpendicular measurements for each particle. Analysis of three independent preparations revealed broad particle-size distributions for both variants (Figure 2C). NC119-3 tended to form larger particles than NC119-2 in two of the three preparations, but this trend was not consistent across batches. Purified nucleocapsid yields were 5.63 ± 1.87 and 6.19 ± 1.33 mg/L culture for NC119-2 and NC119-3, respectively, based on the total mass of the protein-RNA complexes Figure 2D). We next examined whether NC119-3 differed from NC119-2 in the amount and integrity of packaged RNA.

### NC119-3 contained less total RNA while packaging full-length mRNA

We next compared the RNA content of purified NC119-2 and NC119-3 nucleocapsids. Protein and nucleotide concentrations were estimated from absorbance measurements at 260 and 280 nm using extinction coefficient-based equations. ^17^ RNA contents were expressed per 180-subunit equivalent to provide a common protein-based denominator for comparison between the heterogeneous preparations.

NC119-2 and NC119-3 contained (2.20 ± 0.53) × 10³ and (1.11 ± 0.08) × 10³ nucleotides of total RNA per 180-subunit equivalent, respectively (Figure 2E). Full-length mRNA was quantified by quantitative RT-PCR. NC119-2 and NC119-3 contained 0.26 ± 0.18 and 0.19 ± 0.10 molecules per 180-subunit equivalent, respectively (Figure 2E). The mean full-length mRNA content was lower in NC119-3, but he proportional decrease was smaller than that in total RNA. This comparison suggests a greater elative contribution of the encoding mRNA to the total RNA recovered from NC119-3. Equal amounts of total RNA extracted from NC119-2 and NC119-3 were analyzed by denaturing urea-PAGE. RNA species migrating near the expected position of the full-length 863-nt transcript were detected in both samples, together with shorter RNA species (Figure 2F). The band near the expected position of the full-length RNA was more intense in NC119-3 than in NC119-2, consistent with a greater proportion of RNA of this length. Together with the quantitative RT-PCR results, this observation suggests relative enrichment of the encoding mRNA in NC119-3 despite its lower total RNA content.

We next examined RNA protection and particle persistence after treatment with Benzonase (60 kDa) or RNase A (14 kDa). Denaturing urea-PAGE showed that both NC119-2 and NC119-3 retained detectable RNA bands after Benzonase treatment, whereas no RNA bands were detected after RNase A treatment (Figure 2F). A similar difference in nuclease sensitivity was previously reported for NC-1 and NC-2, which contain keyhole-shaped pores. ^13, 15^ DFHBI-1T staining detected RNA species retaining a functional Broccoli aptamer within the BoxB-Br cassette after Benzonase treatment. ^13, 18^ SEC and nsTEM showed that nucleocapsid particles remained intact after Benzonase treatment (Figure 2G and S7). Under the conditions tested, the packaged RNA was more susceptible to RNase A than to Benzonase. This differential susceptibility may reflect differences in nuclease access to the packaged RNA, although it does not by itself establish the presence or size of pores. NC119-2 and -3 cages may possess pores of similar size to those characterized in NC-1 and NC-2 assemblies. ^13, 15^

Following RNase A treatment, nsTEM showed fewer cage-like particles (Figure S7). SEC analysis of RNase A-treated NC119-2 and NC119-3 showed the peak corresponding to fragmented RNAs at ∼11 mL elution fraction (Figure 2G). The cage-sized peak was substantially diminished for NC119-2. NC119-3 showed a small peak at approximately 7–8 mL, indicating that cage-sized species remained after RNase A treatment. These results indicate that some NC119-3 assemblies persisted despite extensive loss of detectable RNA from the sample. However, SEC and urea-PAGE analysis of the bulk sample did not establish whether the remaining assemblies contained RNA. These assemblies may have been RNA-free or retained RNA below the detection limit of the bulk assay. The ability of NC119-3 to package its encoding mRNA, together with the persistence of some cages after RNase A treatment, ed us to conclude the evolution campaign and investigate the structural organization of these assemblies.

### The 180-subunit cage contains position-dependent local structural features

We next used cryo-electron microscopy (cryo-EM) to characterize the architectures within the heterogeneous NC119-3 population. Three-dimensional classification resolved distinct cage families containing 180 and 210 subunits, corresponding to 36 and 42 pentamers, respectively (Figure 3–5 and Figure S8–11). These two cage families accounted for approximately 11% and 12%, respectively, of he particles retained after initial 2D classification, in which all classes containing recognizable particles were selected. Within the 180-subunit family, we first analyzed a tetrahedrally symmetric architecture, designated 180-Tet. The reconstruction reached an overall resolution of 2.90 Å and supported atomic modeling (Figure 3A, S8 and S9). The cage contains 36 pentamers arranged as 12 symmetry-equivalent units, each composed of three pentamers. This overall architecture resembles the 36-pentamer tetrahedral cages previously reported for NC-2 ^15^ and the negatively charged AaLS variant AaLS-neg. ^5^

**Figure 3.**
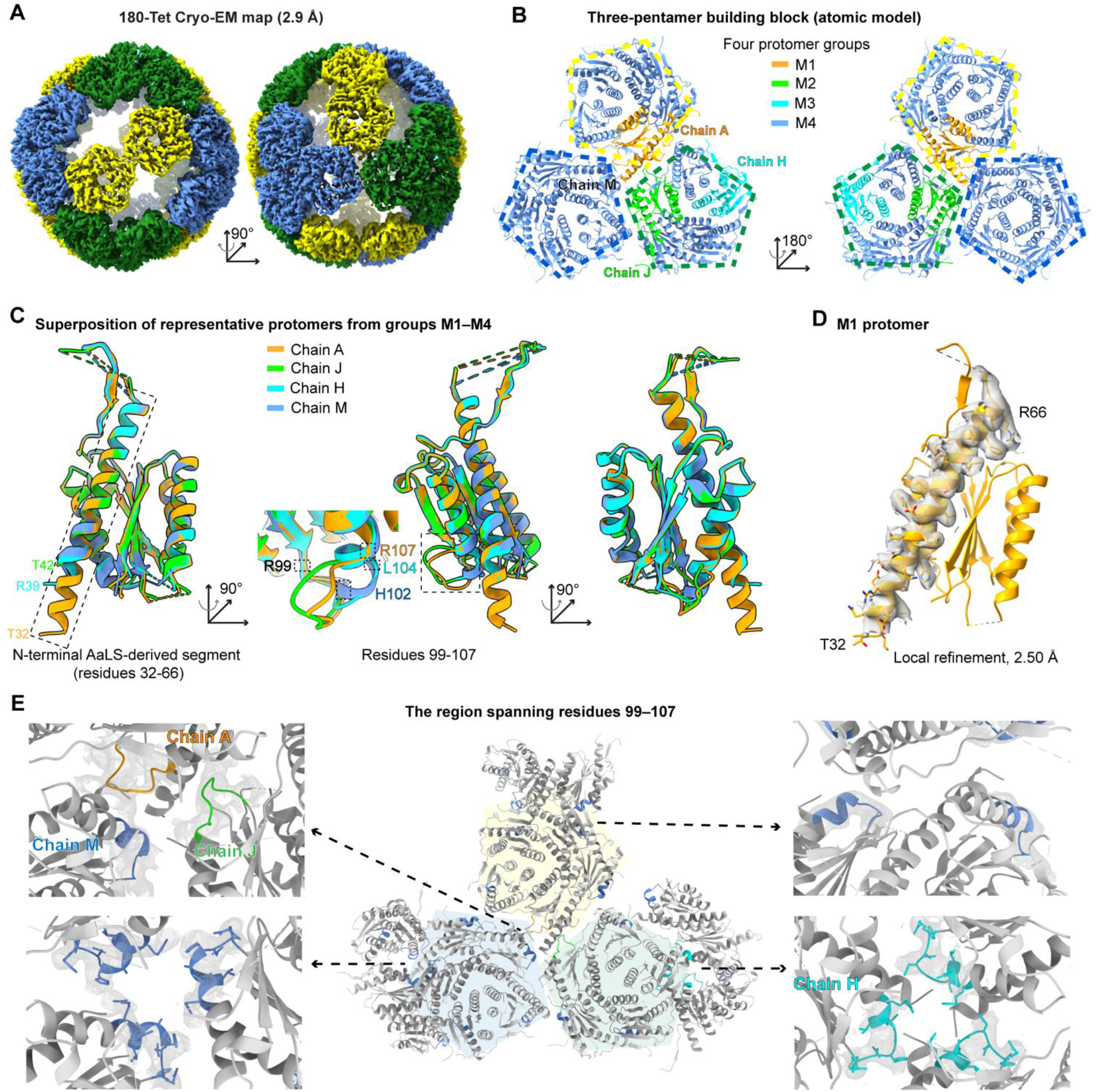
Position-dependent local structural features in the 180-Tet assembly. **A)** Cryo-EM map (EMD-83973) at 2.90 Å in two views related by a 90° rotation. Yellow, green, and blue distinguish three non-equivalent pentamer positions. **(B)** Three-pentamer atomic model (PDB ID: 46PQ) in two views related by a 180° rotation. M1–M4 protomers are orange, lime, cyan, and light blue, respectively, showing their positions within the unit. Dashed pentagon outlines are colored according o the non-equivalent pentamer positions in panel A, independently of the M1–M4 protomer colors. **(C)** Superposition of representative M1-M4 protomers (chains A, J, H, and M), grouped by the resolved *N*-Terminal extension and local loop–helix organization. M1 displays a continuous *N*-terminal helix spanning residues 32–66. Around residues 99–107, helices begin near residue 107 in M1/M2, 104 in M3, and 102 in M4. Groups denote shared local features rather than identical overall conformations. Residue numbering follows the NC119 construct. **(D)** M1 model with density around T32–R66 from the 2.50 Å three-pentamer local reconstruction obtained using symmetry-expanded particles. **(E)** Overview and enlarged views of residues 99–107 in three-pentamer unit. Translucent yellow, blue and green pentagon backgrounds in the central overview indicate the pentamer positions defined in panel A. Residues 99–107 are colored by M1–M4 group as in panel B; the remaining structure is gray. Density s from the original 2.90 Å cryoSPARC map without EMReady2 post-processing.

**Figure 4.**
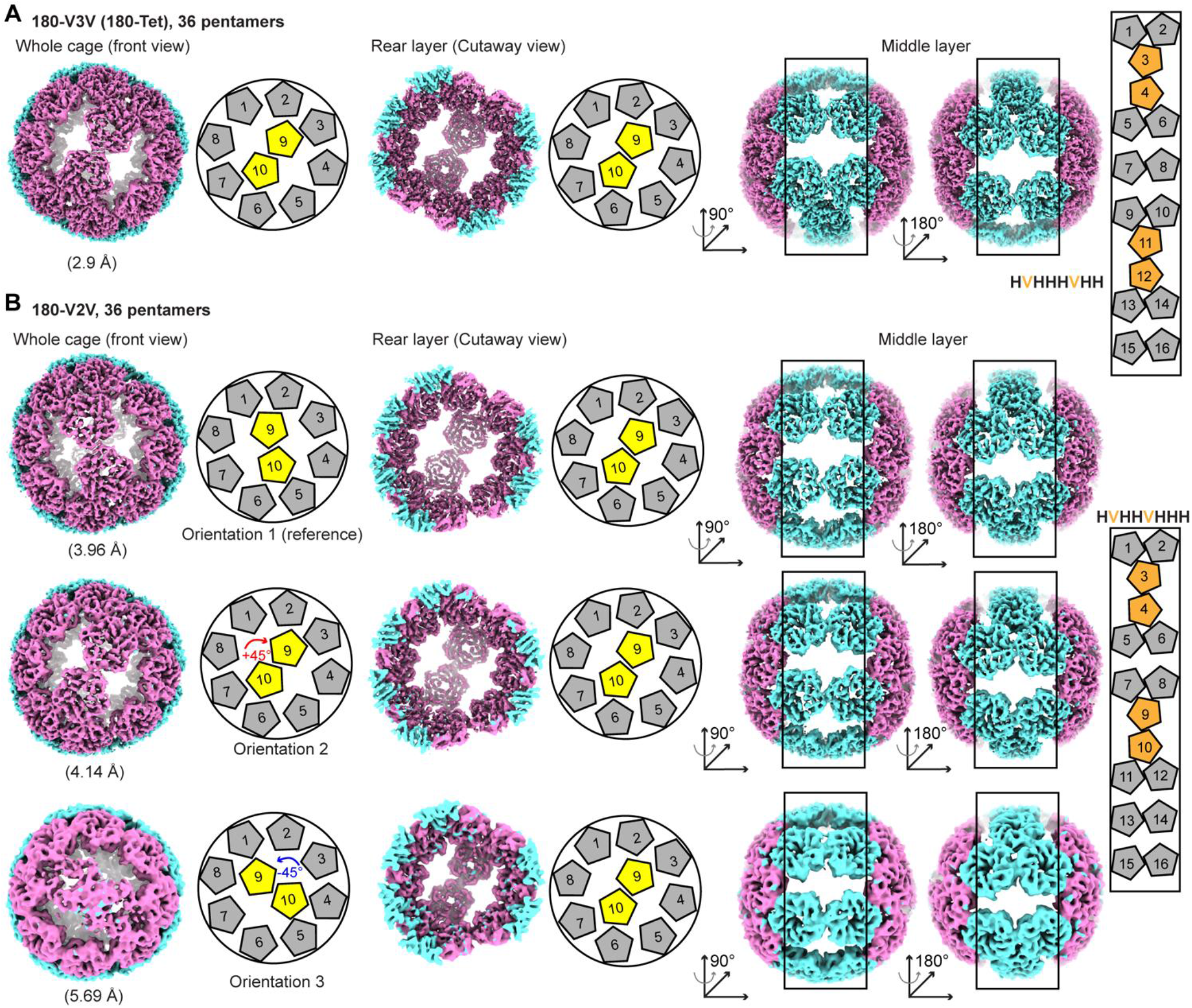
Bridge patterns and terminal orientations of the 180-subunit family. **A)** Cryo-EM reconstruction of the 36-pentamer 180-V3V (180-Tet) architecture at 2.90 Å, shown as whole-cage, cutaway, and middle-layer views with corresponding pentamer schematics. The middle-layer bridge pattern is indicated as HVHHHVHH. **(B)** Cryo-EM reconstructions of three orientations O1–O3) of the 36-pentamer 180-V2V architecture, shown as whole-cage, cutaway, and middle-layer views with corresponding pentamer schematics. The middle-layer bridge pattern is indicated as HVHHVHHH. Map resolutions are indicated in the panels. The front and rear layers are colored in magenta, and the middle layer in cyan. H and V denote horizontal and vertical orientations of paired-pentamer bridges in the reference view. Yellow pentagons highlight the central pentamer pair in each terminal layer; orange pentagons indicate V-oriented bridges. With the rear terminal layers aligned, O1–O3 show front central pentamer-pair orientations corresponding approximately to 0°, +45°, and −45° relative to O1. These angles describe orientations in the geometric schematics rather than directly measured rotations.

**Figure 5.**
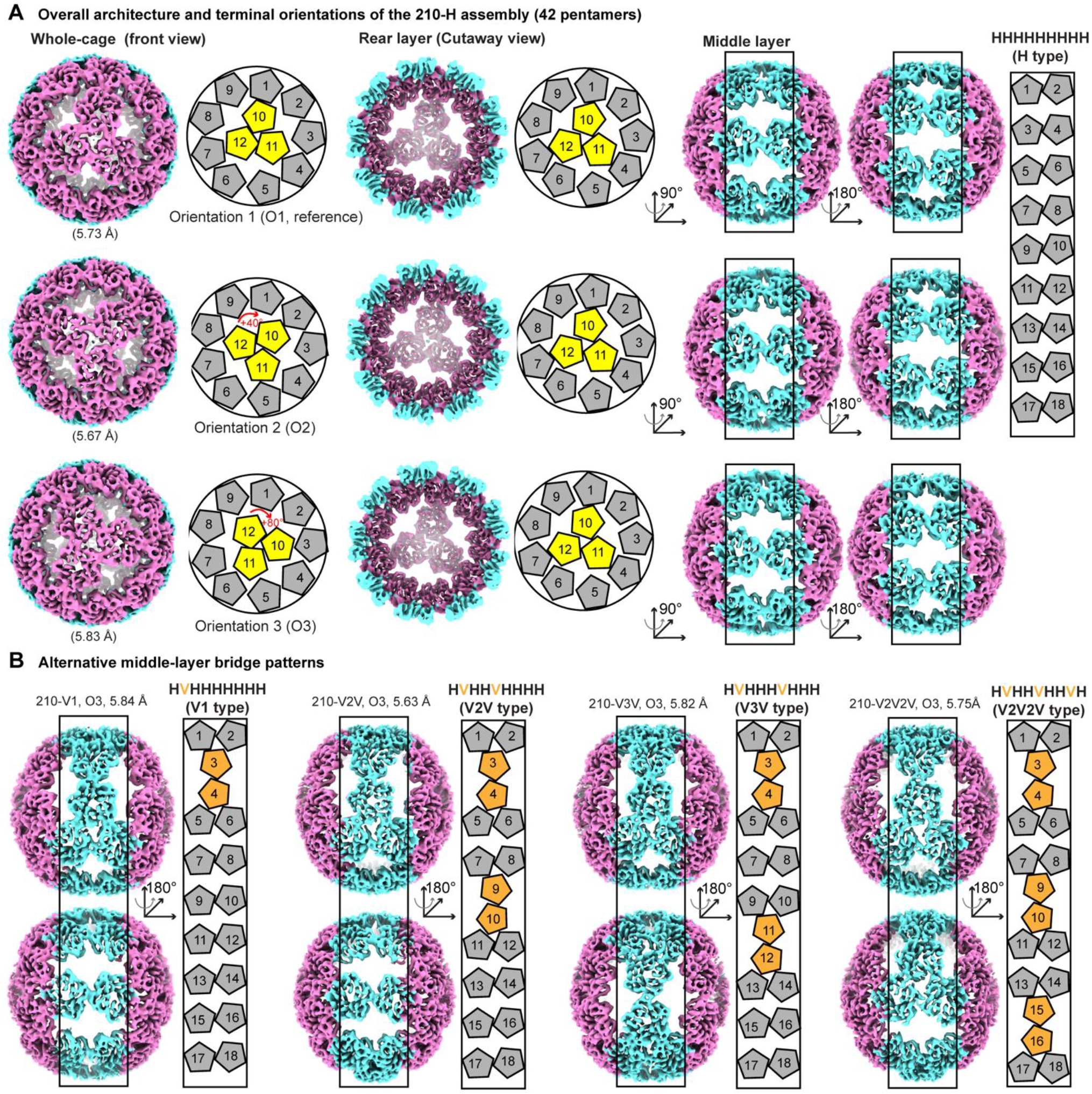
Middle-layer bridge patterns and terminal-orientation space of the 210-subunit family. **A)** Cryo-EM reconstructions of three terminal orientations (O1–O3) of the 42-pentamer 210-H architecture, shown as whole-cage, cutaway, and middle-layer views with corresponding pentamer schematics. The middle-layer bridge pattern is HHHHHHHHH. **(B)** Middle-layer views and corresponding pentamer schematics of the 210-V1, 210-V2V, 210-V3V, and 210-V2V2V architectures. For 210-V1, 210-V2V, and 210-V3V, the displayed O3 maps represent the O3 front-terminal orientation, while the rear-terminal orientations remain mixed. The rear-terminal orientation of 210-V2V2V is resolved. Their bridge patterns are indicated in the panels. Map resolutions are shown for each reconstruction. The front and rear layers are colored in magenta, and the middle layer in cyan.

Within each three-pentamer unit, sequence-identical protomers occupy different local environments. Visual inspection of the cryo-EM density together with comparison of the corresponding atomic models identified four protomer groups, designated M1–M4, based on two local structural features: the extent of the resolved *N*-terminal helical extension and the organization of the loop–helix transition around residues 99–107 (Figure 3B–E). M1 was distinguished by a continuous *N*-terminal helical extension, whereas an equivalent continuous *N*-terminal helical extension was not resolved in the other protomers. The remaining protomers were divided into M2–M4 according to the loop–helix transition, with helices beginning near residues 107, 104, and 102, respectively (Figure 3C). M1 shared the loop–helix organization of M2. Residue numbers in this section refer to the NC119 construct unless otherwise indicated and differ from those of AaLS-wt because of circular permutation and the addition of the λN+ peptide and linker.

The M1 model contained an extended *N*-terminal helical segment beginning at T32. This residue corresponds to T120 of AaLS-wt and marks the beginning of the AaLS-derived sequence following the λN+ peptide and linker. To examine this region more closely, we performed local refinement of a three-pentamer unit using the symmetry-expanded particle set. Aligning and refining individual three-pentamer units separately can reduce blurring caused by possible relative movements between units hat remain comparatively rigid internally. The resulting 2.50 Å map supported a continuous helical backbone from T32 to R66 in M1 (Figure 3D and S8). Spatial constraints near the pseudo-threefold center of the three-pentamer unit may prevent all three *N*-terminal segments from simultaneously adopting the extended helical conformation observed in M1.

The local loop–helix differences further distinguished M2–M4 (Figure 3C, E). In M2, residues 99–106 formed a loop followed by an α-helix beginning at residue 107, with the modeled backbone agreeing with the corresponding density. In M3, helical density was apparent from approximately residue 104, although the precise backbone conformation within residues 100–104 remained uncertain. In M4, residues 99–101 formed a loop followed by a continuous α-helix beginning near residue 102, with both regions supported by the density. M4 protomers shared this local organization despite differences in their modeled *N*-terminal lengths and interprotomer contacts; this grouping does not imply identical conformations throughout the protomers.

The density-supported differences in residues 99–107 occurred in distinct local contact environments involving neighboring pentamers (Figure 3E). Together with the extended *N*-terminal helix resolved in M1, these observations reveal position-dependent variation in local structure and ordering within the 180-Tet cage. We next compared the architectures of the 180-subunit cages to examine variation in bridge arrangement and the orientation of terminal pentamer-pair modules.

### The 180-subunit cages differ in bridge arrangement and terminal pentamer-pair module orientation

We next compared the pentamer arrangements within the 180-subunit family. In addition to 180-Tet, three-dimensional classification identified three additional classes with geometrically distinct arrangements, each containing 36 pentamers (Figure 4). For comparison, we partitioned each cage into two terminal layers (front and rear), each containing ten pentamers, and a middle layer containing sixteen pentamers. Each terminal layer consists of a peripheral ring of eight pentamers surrounding a central pentamer-pair module (Figure 4A, B, left). The middle layer comprises eight pentamer pairs that bridge the two terminal layers and form a ring around the cage. These paired-pentamer units adopt two orientations, designated horizontal (H) and vertical (V) in the reference view used for comparison Figure 4A, B, right).

Two middle-layer bridge patterns were identified. The 180-Tet architecture contains six H-oriented and wo V-oriented bridges arranged in the sequence “HVHHHVHH” (Figure 4A, right). The two V-oriented bridges are separated by three H-oriented bridges on either side of the ring, and this arrangement is designated 180-V3V. A second bridge pattern, designated 180-V2V, has the sequence “HVHHVHHH” Figure 4B, right). In this arrangement, the two V-oriented bridges are separated by two H-oriented bridges on one side and four on the other. The “VnV” designation uses the smaller number of intervening H-oriented bridges. Both patterns contain the same numbers of H- and V-oriented bridges but differ in the placement of the V-oriented bridges around the cage.

The 180-V2V population was further separated into three classes, designated O1, O2, and O3. These classes share the same middle-layer bridge pattern but differ in the orientation of the central pentamer pair in the front terminal layer (Figure 4B, left). With the rear terminal layer aligned to a common reference, the two central pentamers at the front occupied three discrete orientations relative to the surrounding eight-pentamer ring. In the geometric model, these orientations correspond approximately o 0°, +45°, and −45° relative to O1. A fourth orientation at approximately +90° is geometrically possible, but a corresponding class was not identified (Figure S10A). This suggests the relative positioning between the central pentamer pairs and V-form bridges are not random, but they have preference.

The 180-subunit cages exhibit multiple geometric arrangements through variation in both middle-layer bridge patterns and terminal pentamer-pair orientations. Although the geometric model permits four terminal orientations for each of the V3V and V2V bridge patterns, the classified structures comprised one V3V arrangement and three V2V arrangements. This selective recovery suggests that bridge arrangement and terminal orientation may be associated, with certain combinations favored over others. We next examined how these architectural features vary in the larger 210-subunit cages.

### The 210-subunit family combines multiple bridge patterns with alternative terminal-module orientations

We next examined the 210-subunit family, in which each cage contains 42 pentamers. For comparison, we partitioned each cage into two terminal layers containing twelve pentamers each and a middle layer containing eighteen pentamers (Figure 5A). Each terminal layer consists of a peripheral ring of nine pentamers surrounding a central three-pentamer module. The middle layer comprises nine paired-pentamer bridges connecting the terminal layers and arranged in a ring, one more than in the 180-subunit family. The H and V orientations defined for the 180-subunit bridges were also used to describe these arrangements.

Classification identified five middle-layer bridge patterns, designated 210-H, 210-V1, 210-V2V, 210-V3V, and 210-V2V2V (Figure 5 and S11). The 210-H architecture contains only H-oriented bridges, and 210-V1 contains a single V-oriented bridge. The 210-V2V and 210-V3V architectures each contain wo V-oriented bridges but differ in their spacing around the ring, with a minimum of two and three intervening H-oriented bridges, respectively. The 210-V2V2V architecture contains three V-oriented bridges, each separated from the next by two H-oriented bridges. These five patterns preserve the total number of pentamers while varying the number and positions of V-oriented bridges.

Similar to 180-mer cage’s case, variation in terminal-module orientation was observed within the 210-mer cages. The central three pentamers within each terminal layer occupied 3 distinct orientations. The 210-H and 210-V2V2V architectures were compatible with C3 symmetry and were refined accordingly Figure S11 and S12). With the rear terminal layer aligned to a common reference, the central three-pentamer module at the front occupied three discrete orientations within the surrounding nine-pentamer ring. These classes were designated O1, O2, and O3 (Figure 5A and S10B). In the geometric model, heir orientations correspond approximately to 0°, 40°, and 80° relative to O1.

Unlike the 210-H and 210-V2V2V architectures, the middle-layer bridge patterns of 210-V1, 210-V2V, and 210-V3V do not preserve global C3 symmetry, although the central three-pentamer modules retain local C3 symmetry. These architectures were reconstructed without imposing symmetry (C1) (Figure S11). Within each bridge-pattern class, classification resolved three orientations of the front central three-pentamer module relative to the asymmetric middle layer. The corresponding rear module remained heterogeneous, and its orientation subclasses were not further classified. The geometric model permits three candidate orientations at each end, yielding up to nine possible combinations of front and rear terminal-module orientations (Figure S10C).

Together, the classified structures reveal variation in both the number and spacing of V-oriented bridges and the orientation of terminal three-pentamer modules within the 42-pentamer cages. These findings extend the architectural diversity observed in the 180-subunit family to larger cages while retaining the organization into terminal layers connected by paired-pentamer bridges.

### NC119-3 exhibits structural variation within and between cages

In addition to the 180- and 210-mer cages, classification identified a minor population consistent with an approximately 190-subunit cage containing 38 pentamers (Figures S8 and S13). Two related reconstructions were obtained at 8.29 Å and 8.42 Å resolution (Figure S13). The maps were consistent with two terminal layers containing ten pentamers each and a middle layer containing eighteen pentamers. The proposed middle-layer arrangements included three-pentamer bridges alongside paired-pentamer bridges, suggesting that bridge composition may provide an additional source of architectural variation.

The better-resolved 180- and 210-subunit structures reveal structural variation at several organizational levels. Within 180-Tet (V3V), sequence-identical protomers exhibit position-dependent differences in local conformation and *N*-terminal ordering (Figure 3B–E). Extended helical densities resembling the *N*-terminal helix of the M1 protomer were also observed at three-pentamer junctions across multiple NC119-3 cage architectures, with representative examples shown for 180-V2V O1 and 210-V2V O3 Figure S14A, B). These observations suggest that local environments supporting *N*-terminal ordering may be shared across distinct cage architectures. Between cages of the same stoichiometry, structural differences involve alternative bridge arrangements and orientations of the central pentamer pair Figure 4) or three-pentamer module (Figure 5) within the terminal layers. Variation between the cage families additionally involves differences in the total number of pentamers. The tentative 190-subunit reconstructions further suggest that variation in the number of pentamers forming individual bridges may provide another means of diversifying cage architecture.

Beyond these reconstructed architectures, larger cage-like particles were also observed in the 2D class averages (Figure S8), but did not yield interpretable three-dimensional reconstructions. Structural heterogeneity may have contributed to the difficulty in resolving these particles. These observations suggest that the resolved structures capture only part of the architectural diversity of NC119-3, with larger particles potentially accommodating additional pentamer arrangements.

Evolution of cpAaLS119 yielded NC119-3 with several pentamer-based cage architectures. These architectures differ from the domain-swapped *T* = 4 architecture previously characterized in cpAaLS84-derived NC-3 and NC-4. ^15^ The two evolution campaigns also differed in selection history and clone choice. Selection from cpAaLS84-derived NC-2 to NC-3 used RNase A in place of Benzonase, ^13^ whereas NC119 was selected with only Benzonase. In the NC119 campaign, we deliberately advanced he morphologically heterogeneous NC119-2 variant during clone screening. Comparative evolution experiments under matched selection and screening conditions would help distinguish the contribution of backbone connectivity from those of the evolutionary protocols.

The parental cpAaLS119 scaffold was already structurally polymorphic. ^12, 16^ Some architectures resolved in NC119-3 may therefore have been present before evolution but occurred at low abundance or were not resolved in earlier analyses. Sequence changes could have altered the relative abundance or stability of these states. Comparable structural analysis of the parental scaffold and intermediate variants would help distinguish these changes from the appearance of new architectures during evolution.

RNA cargo is another variable that may influence the diversity of NC119-3 cage architectures. NC119 construct contains the λN+ peptide and recruits its encoding RNA during intracellular assembly. At lower contour levels, diffuse internal density was visible in multiple cage reconstructions calculated without imposed symmetry, with representative examples shown for 180-V2V O1 and 210-V2V O3 Figure S14C, D). This density may correspond in part to encapsulated RNA, although it was insufficiently resolved for unambiguous assignment and may also include contributions from unresolved protein segments. Bulk RNA analysis detected mRNA together with shorter RNA species (Figure 2E, F), but the cargoes of individual particles and structural classes were not determined. Particles formed by the same protein sequence may therefore contain different RNA cargoes. Differences in RNA length, sequence, secondary structure, or occupancy could favor particular pentamer arrangements or cage stoichiometries. ^19, 20^

Related systems provide examples of cargo-dependent changes in nucleocapsid morphology and local protein conformation. DNA origami templates have been used to direct the size and shape of both the artificial nonviral nucleocapsid spNC-4 ^21^ and assemblies formed by viral capsid proteins. ^22^ In bacteriophage MS2, binding of an RNA stem-loop alters coat-protein dimer conformation and regulates capsid assembly. ^23–25^ Establishing an *in vitro* reconstitution system using purified NC119 proteins and defined RNA cargoes would enable direct tests of how cargo affects cage stoichiometry, pentamer arrangement, and local protomer conformation.

## Conclusions

Directed evolution of a cpAaLS119-derived progenitor yielded NC119-3, whose protein subunits assembled into structurally heterogeneous, pentamer-based cages that packaged their encoding mRNA. Recovery of the assembled nucleocapsids was improved relative to the progenitor. Cryo-EM analysis resolved 180- and 210-subunit cage families and identified a minor population consistent with an approximately 190-subunit architecture. Larger cage-like particles exceeding the dimensions of the resolved 210-subunit cages were also observed in 2D class averages but did not yield interpretable three-dimensional reconstructions. Within the 180- and 210-subunit families, cages with the same subunit stoichiometry differed in bridge arrangement and in the orientation of the central pentamer pair or central three-pentamer module, respectively. Within 180-Tet, one architecture in the 180-subunit family, sequence-identical protomers exhibited position-dependent differences in local conformation and *N*-terminal ordering. The pentamer-based architectures observed in NC119-3 differ from the domain-swapped *T* = 4 architecture characterized in the cpAaLS84 lineage. Parallel evolution experiments under matched selection and screening conditions would help clarify the contribution of backbone connectivity to this difference. Together, this study shows that functional evolution from distinct topological starting points can yield RNA-packaging nucleocapsids with different cage architectures.

## Experimental Section

### Expression and Purification of Capsids

NC119 capsids were produced in *E. coli* BL21(DE3)-Gold (Agilent Technologies) transformed with pMG-dB vectors encoding the corresponding NC119 variants or libraries. The pMG-dB vector contains BoxB-Br tags in both the 5′ and 3′ UTRs of the NC119-encoding transcript. The cells were grown at 37°C in Luria Bertani (LB) medium until the OD600 reached 0.4–0.6, at which point protein production was induced by adding isopropyl β-d-1-thiogalactopyranoside (IPTG) to a final concentration of 0.3 mM. Then, the cells were cultured at 37°C for 5 h. The cells were harvested by centrifugation at 5,000 × g and 4°C for 10 min. The cell pellet was frozen by liquid nitrogen and stored at −80°C until purification. The cell pellet from a 1-L culture was resuspended in 10 mL lysis buffer [50 mM sodium phosphate buffer (pH 7.4) containing 1 M NaCl and 20 mM imidazole]. After lysis by sonication and clearance by centrifugation at 12,000 × g and 25°C for 10 min, the supernatant was loaded onto 1 mL of Nuvia™ MAC Resin (Ni-charged-NTA, Bio-Rad) in a gravity flow column. After washing with lysis buffer, protein capsid was eluted with 2.5 mL elution buffer [50 mM sodium phosphate buffer (pH 7.4) containing 300 mM NaCl and 500 mM imidazole]. The eluted solution was mixed with EDTA to a final concentration of 5 mM. Protein capsids were further purified by size-exclusion chromatography (SEC) [column, TSKgel G5000PWxl (Tosoh Bioscience); eluent, 50 mM sodium phosphate buffer (pH 7.4) containing 200 mM NaCl and 5 mM EDTA; flow rate, 1 mL/min; room temperature]. Protein and RNA concentrations were determined by UV absorbance following a previously reported protocol. ^17^ Extinction coefficients for proteins were calculated using the ExPASy ProtParam tool (https://web.expasy.org/protparam/).

### Nuclease Treatment and RNA Extraction from Purified Capsids

After size-exclusion chromatography, the protein capsids containing ∼3000 ng RNA were mixed with 2 μL Benzonase (70746-4CN, 25 U/μL; Merck Millipore) or 0.2 μL RNase A (EN0531, 10 mg/mL; Thermo Fisher Scientific) at 37 °C for 1 h in 50 mM sodium phosphate buffer (pH 7.4) containing 200 mM NaCl, 5 mM EDTA, and 10 mM MgCl_2_. Following nuclease treatment, RNAs were extracted with TRIzol reagent (15596026; Invitrogen) and dissolved in water.

### epPCR Library Construction

epPCR was carried out by using the JBS Error-Prone Kit (PP-102; Jena Bioscience) according to the manufacturer’s protocols. Primers priHL0024(F) and priHL0027(R) (Table S3) and template pMG-dB-NC119-1 were used for the library in round 1, and primers priHL0032(F) and priHL0033(R) and template pMG-dB-NC119-2 were used for the library in round 2. Purified epPCR products and the acceptor vector were digested with NdeI/XhoI (New England Biolabs) for pMG-dB-NC119-1 and pMG-dB-NC119-2. The fragments were ligated with T4 DNA ligase (LGK-201; TOYOBO) and purified. Electrocompetent E. coli HST08 Premium Electro-Cells (9028; Takara) were transformed with the capsid libraries approximately 700 ng ligation product) by electroporation and then incubated in 3 mL LB medium for 1 h at 37°C. The transformation library size was estimated by plating serial dilutions of the cells onto LB agar containing ampicillin (100 μg/mL), yielding approximately 1–4 × 10⁶ transformants per round of directed evolution. The remaining cells were cultured overnight at 37°C and 180 rpm. The next day, plasmid DNA was extracted using the FastGene™ PlasmidMini Kit (FG-90502, Nippon Genetics).

### Round 1 of Directed Evolution

The ORF of pACTet-N+(GA)4-cpAaLS119 was amplified by PCR using primers priHL0024(F) and priHL0005(R) (Table S3). The purified PCR product and acceptor vector pMG-dB-λN+cpAaLS84 were digested with NdeI and XhoI, purified, and ligated to generate pMG-dB-λN+cpAaLS119 (pMG-dB-NC119-1). An epPCR library was then prepared as described in epPCR Library Construction.

Electrocompetent E. coli BL21(DE3) cells (DS258; BioDynamics) were transformed with approximately 1 μg of pMG-dB-NC119-1-Lib1 by electroporation and recovered in 3 mL LB medium for 1 h at 37 °C. The culture was transferred to 22 mL LB medium containing ampicillin (100 μg/mL) and grown overnight at 37 °C and 180 rpm. The preculture was then transferred to 1 L LB medium and grown to an OD600 of 0.4–0.6. Protein expression was induced with 0.3 mM IPTG for 5 h at 37 °C and 160 rpm.

Cells were harvested by centrifugation at 5,000 × g for 10 min at 4 °C and resuspended in 10 mL lysis buffer containing 50 mM sodium phosphate (pH 7.4), 1 M NaCl, and 20 mM imidazole. Following sonication, the lysate was clarified by centrifugation at 12,000 × g for 10 min at 25 °C. The supernatant was loaded onto 1 mL Nuvia IMAC resin (Ni-charged NTA; Bio-Rad) in a gravity-flow column. After washing with lysis buffer, nucleocapsids were eluted with 2 mL of 50 mM sodium phosphate (pH 7.4), 300 mM NaCl, and 500 mM imidazole. EDTA was added to the eluate to a final concentration of 5 mM. Nucleocapsids were further purified by SEC using a TSKgel G5000PWxl column (Tosoh Bioscience) equilibrated with 50 mM sodium phosphate (pH 7.4), 200 mM NaCl, and 5 mM EDTA at a flow rate of 1 mL/min at room temperature.

### First cycle of Benzonase selection

IMAC-purified nucleocapsids containing approximately 2500 ng of nucleic acids were treated with 0.2 μL Benzonase (70746-4CN, 25 U/μL; Merck Millipore) for 1 h at 37°C in IMAC elution buffer supplemented with 10 mM MgCl₂. Protected RNA was extracted using TRIzol reagent (15596026; Invitrogen) and dissolved in water. First-strand cDNA was synthesized using SuperScript IV Reverse Transcriptase (18090200; Invitrogen) and primer priHL0027(R) according to he manufacturer’s protocol. The resulting cDNA was amplified by step-down PCR using KOD One DNA polymerase (KMM-201X5; TOYOBO) and primers priHL0024(F) and priHL0027(R) (Table S3). The amplified DNA was purified, digested, and ligated into the pMG-dB vector to generate NC119-1-Lib2.

### Second and third cycles of Benzonase selection

Electrocompetent *E. coli* BL21(DE3) cells were transformed with the library obtained from the preceding selection cycle. Following expression and purification as described above, SEC-purified nucleocapsids containing approximately 1200 ng of nucleic acids were treated with 0.2 μL Benzonase for 1 h at 37 °C in SEC buffer supplemented with 10 mM MgCl₂. Protected RNA was extracted and subjected to reverse transcription and PCR as described above. The amplified DNA was purified, digested, and ligated into the pMG-dB vector to generate the library for the subsequent cycle. This procedure generated NC119-1-Lib3 and subsequently NC119-1-Lib4.

Following three cycles of Benzonase selection, 10 individual clones were randomly selected from NC119-1-Lib4 and analyzed by sequencing, expression, purification, and nsTEM.

### Round 2 of Directed Evolution

An epPCR library was prepared from pMG-dB-NC119-2 as described in epPCR Library Construction. Electrocompetent *E. coli* HST08 Premium Electro-Cells (9028; Takara) were transformed with approximately 700 ng of the resulting ligation product, NC119-2-Lib1, by electroporation. Library propagation, protein expression, and nucleocapsid purification were performed as described for Round 1.

SEC-purified nucleocapsids containing approximately 3000 ng of RNA were treated with 2 μL Benzonase (70746-4CN, 25 U/μL; Merck Millipore) for 1 h at 37 °C in SEC buffer supplemented with 10 mM MgCl₂. Protected RNA was extracted and subjected to reverse transcription and PCR as described for Round 1. The amplified DNA was purified, digested, and ligated into the pMG-dB vector o generate the library for the subsequent selection cycle. Three cycles of Benzonase selection were performed.

Following three cycles of Benzonase selection, 25 individual clones were randomly selected from NC119-2-Lib4 and analyzed by sequencing, expression, IMAC purification, and nsTEM.

### Denaturing PAGE Analysis

RNA samples were mixed with an equal volume of 2× RNA loading buffer containing 10 M urea, 2 mM EDTA, and 2 mM Tris-HCl (pH 7.5), heated at 95 °C for 5 min, and then chilled on ice. Denatured RNA samples were separated on 6% polyacrylamide gels containing 8 M urea in 1× TBE. DynaMarker RNA Low II (MK-L; 20–500 nt; DM152) and DynaMarker RNA High (MK-H; 200–8,000 nt; DM160), both from BioDynamics Laboratory Inc., were used as RNA size markers. To visualize BoxB-Br-containing RNA, he gels were first stained with DFHBI-1T (410; Lucerna) and imaged. The gels were then washed to remove DFHBI-1T and subsequently stained with EzPreStain DNA&RNA (WSE-7135; ATTO) to visualize total RNA, following a previously reported protocol. ^18^ Fluorescence signals were acquired using a Typhoon FLA 9500 laser scanner (GE Healthcare) with excitation at 473 nm. The same imaging settings were used for DFHBI-1T and EzPreStain detection.

### Quantification of Encapsulated RNA

The total nucleic acid content of NC119 capsids was determined by UV absorbance measurements according to a previously reported method. ^17^ The nucleic acid content was calculated as the number of nucleotides per subunit and expressed as nucleotides per 180-mer.

To quantify encapsulated full-length mRNA, quantitative RT-PCR (qRT-PCR) was performed using RNA extracted from purified NC119 capsids. RNA concentrations were measured using a Qubit RNA BR Assay Kit (Q10210; Invitrogen). The extracted RNA was diluted to 80 ng/μL, and 2 μL (160 ng) was reverse-transcribed into first-strand cDNA using SuperScript IV Reverse Transcriptase (18090200; nvitrogen) with the priHL0005(R) primer according to the manufacturer’s instructions. For qPCR, the resulting cDNA was diluted 1,000-fold, and 1 μL of the diluted cDNA was mixed with PowerUp SYBR Green Master Mix (A25779; Applied Biosystems) and 0.4 μL each of 10 μM priHL0050(F) and priHL0027(R) primer (Table S3) stocks. The gene-specific reverse transcription primer was located downstream of the 3′ BoxB-Br cassette, whereas the qPCR primers amplified a 739-bp region extending rom upstream of the 5′ BoxB-Br cassette to upstream of the 3′ BoxB-Br cassette. Real-time PCR was performed using a StepOnePlus Real-Time PCR System (4376600; Applied Biosystems). Standard curves were generated using serial dilutions of in vitro-transcribed NC119 mRNA of known concentration. A no-reverse-transcriptase (−RT) control was included to assess DNA contamination, and no amplification was detected.

### Calculation of Full-Length mRNA Copies per 180-mer

The number of full-length mRNA copies per 180-mer was calculated in three steps. First, qRT-PCR was used to determine the number of full-length mRNA copies per nanogram of total extracted RNA. Second, the total RNA mass per 180-mer was calculated from the UV absorbance-derived RNA content nt/subunit) by multiplying by 180 and converting nucleotide number to RNA mass using an average nucleotide molecular weight of 340 Da. Finally, the number of full-length mRNA copies per 180-mer was calculated by multiplying the full-length mRNA copies per nanogram of total RNA by the total RNA mass per 180-mer. The 180-mer was used as a standardized normalization basis for comparison between NC119 variants.

### Statistical Analysis

Nucleocapsid yields, total RNA content, and full-length mRNA copy numbers are presented as mean ± SD from three independent culture, expression, and purification experiments per variant (n = 3). Comparisons between NC119-2 and NC119-3 were performed using two-tailed unpaired Welch’s *t*ests in GraphPad Prism version 11.0.0 (93), without assuming equal population variances. *P* values were not adjusted for multiple comparisons. A *p* value < 0.05 was considered statistically significant.

### Negative-Stain Transmission Electron Microscopy

Negative-stain transmission electron microscopy (nsTEM) was performed as reported previously. ^26^ SEC-purified NC119 samples (2 μM monomer) were adsorbed onto glow-discharged, carbon-coated grids for 1 min and blotted. The grids were stained with 2% (w/v) phosphotungstic acid (PTA, pH 7.0) using three successive drops. The first two drops were immediately blotted, whereas the third was left on the grid for 20 s before blotting. The grids were air-dried prior to imaging. TEM images were acquired using a Hitachi H7650 Zero.A transmission electron microscope.

Particle diameters were measured from nsTEM images using Fiji (version 2.1.0/1.53c). ^27^ For each particle, the diameter was calculated as the average of two orthogonal dimensions to account for deviations from a circular projection. All measurable particles from the acquired micrographs were included. Measurements from three independent sample preparations were plotted separately for visualization of the overall particle-size distribution. In total, 138 NC119-2 particles and 126 NC119-3 particles were analyzed. The distributions were visualized as violin plots, with the median and interquartile range indicated.

### Cryo-Electron Microscopy Sample Preparation and Data Collection

Freshly purified NC119-3 was concentrated to 4–5 mg/mL in SEC buffer (50 mM sodium phosphate, pH 7.4; 200 mM NaCl, and 5 mM EDTA). Quantifoil R1.2/1.3 Cu 300 grids were glow-discharged at 7 Pa and 10 mA for 10 s using an Auto Fine Coater (JEC-3000FC, JEOL). A 3 μL aliquot of sample was applied to each grid and blotted for 2 s at a blot force of 10, 100% humidity, and 22 °C using a Vitrobot Mark IV System (Thermo Fisher Scientific). The grids were plunge-frozen in liquid ethane and stored n liquid nitrogen until data collection.

Cryo-EM data were collected at SPring-8 using a CRYO ARM 300 microscope (JEOL) operated at 300 kV and equipped with an in-column Omega energy filter and a K3 direct electron detector (Gatan) operated in correlated double sampling (CDS) electron-counting mode. YoneoLocr ^28^ was used for hole detection, and automated data acquisition was performed using SerialEM ^29^. Images were recorded at a nominal magnification of 60,000×, corresponding to a calibrated pixel size of 0.752 Å. Movies were acquired over 50 frames with a total electron exposure of approximately 50.45 e⁻/Å², using a 5 × 5 beam-image-shift acquisition pattern. The applied defocus was set between −1.4 and −1.6 μm in 0.1 μm increments at each stage shift.

### Cryo-EM Image Processing

Single-particle cryo-EM analysis of NC119-3 was performed using cryoSPARC v4.5.3. ^30^ A total of 18,578 movies were motion-corrected using Patch Motion Correction, followed by contrast transfer unction (CTF) estimation using Patch CTF Estimation in cryoSPARC. An initial three-dimensional cage reconstruction was obtained from a preliminary set of picked particles and used to generate two-dimensional projection templates with the Create Templates job in cryoSPARC. Template-based particle picking using these projections yielded 3,443,815 particles. Following particle extraction and inspection, 1,782,592 particles were subjected to 2D classification into 100 classes, from which 1,532,817 particles were selected for subsequent analysis. Detailed processing workflows and particle numbers for the 180- and 210-subunit assembly families are summarized in Figures S8 and S11.

For the 180-subunit family, particles were separated by heterogeneous refinement in C1 and further defined to resolve distinct assembly states. The 180-V3V population was subjected to homogeneous refinement in C1 and tetrahedral symmetry, followed by reference-based motion correction and a second tetrahedral refinement. A total of 112,851 particles were then symmetry-expanded using the tetrahedral point group, generating 1,354,212 symmetry-expanded particle instances. The expanded particles were subjected to focused 3D classification without alignment into 10 classes in C1, using a filter resolution of 4 Å and a mask encompassing one three-pentamer module. PCA initialization was used for the classification. Selected classes comprising 1,020,460 symmetry-expanded particle instances were combined and reconstructed in C1, yielding a 2.90 Å map. Subsequent local refinement using a mask encompassing the three-pentamer module yielded a 2.50 Å reconstruction. Alternative 180-subunit architectures and the minor approximately 190-subunit population were resolved by further C1 refinement and 3D classification as detailed in Figure S8. Resolution assessments are provided in Figures S9 and S13, and cryo-EM data collection, reconstruction, and model validation statistics are summarized in Figure S15.

Enlarged-particle subsets were further processed to resolve the 210-subunit assembly family. Particles were subjected to successive rounds of 2D classification, heterogeneous refinement, homogeneous refinement, and 3D classification in C1 to separate distinct architectures and terminal-module orientations. The resulting classes were further refined and reconstructed using C1 or C3 symmetry, as appropriate for each processing stage. These analyses resolved the 210-H, 210-V1, 210-V2V, 210-V3V, and 210-V2V2V architectures and their alternative terminal-module orientations. For 210-V1, 210-V2V, and 210-V3V, focused classification resolved distinct front-terminal orientations, while the rear terminal orientations remained heterogeneous. Detailed classification pathways, particle numbers, and reconstruction resolutions are provided in Figure S11, and cryo-EM data collection and reconstruction statistics are summarized in Figure S16. Global resolutions were determined using the gold-standard Fourier shell correlation criterion at FSC = 0.143; resolution assessments for the C3-symmetric 210-H and 210-V2V2V reconstructions are provided in Figure S12.

### Atomic Model Building and Refinement

An initial model of the NC119-3 monomer was predicted using AlphaFold 3 via the AlphaFold Server. ^31^ Atomic model building was initiated using the 2.50 Å locally refined map of the three-pentamer module. To improve map interpretability, the locally refined map was post-processed using EMReady2. ^32^ The AlphaFold 3-predicted monomer model was fitted into the EMReady2-processed map and initially adjusted using the ISOLDE ^33^ plugin in UCSF ChimeraX v1.11 ^34^. The three-pentamer model was subsequently manually built and adjusted in Coot v1.1.20 ^35^, and refined in Phenix v2.1-6048 ^36^ against the EMReady2-processed map. The resulting three-pentamer model was propagated by non-crystallographic symmetry (NCS) to generate an initial full-cage model. The full-cage model was subsequently refined in Phenix against an EMReady2-processed version of the 2.90 Å 180-V3V reconstruction. Following further manual adjustment of the three-pentamer model, NCS was reapplied o generate the final full-cage atomic model. Final map–model validation was performed against the original 2.90 Å cryoSPARC reconstruction without EMReady2 post-processing. Model validation statistics are summarized in Figure S15.

## Associated Content

Supporting Information: Supplementary sequence information, biochemical characterization, negative-stain TEM analyses, and detailed cryo-EM image-processing workflows and structural analyses of the NC119 assemblies (PDF and Excel files).

## Data Availability

Cryo-EM density maps of the NC119-3 artificial nucleocapsids have been deposited in the Electron Microscopy Data Bank (EMDB) under accession codes EMD-83973, EMD-84003, EMD-84005–84008, EMD-84017, EMD-84019, EMD-84020, EMD-84103, EMD-84104, EMD-84112, EMD-84116, EMD-84118, EMD-84121, EMD-84124, EMD-84126, EMD-84127, and EMD-84130–84133. The atomic coordinates of the 180-V3V (180-Tet) nucleocapsid have been deposited in the Protein Data Bank PDB) under accession code 46PQ. The correspondence between maps and structural classes is provided in Figures S15 and S16. Additional data supporting the findings of this study are provided in he Supporting Information or are available from the corresponding author upon reasonable request.

## Author Information

The authors declare no competing financial interest.

## Acknowledgments

Cryo-EM grids were screened using EM02CT, and cryo-EM data were collected using EM01CT at SPring-8 (Hyogo, Japan), with the approval of the Japan Synchrotron Radiation Research Institute JASRI) under proposal number 2025B2532 to T.N. This study was partially supported by the Research Support Project for Life Science and Drug Discovery (Basis for Supporting Innovative Drug Discovery and Life Science Research (BINDS)) from Japan Agency for Medical Research and Development AMED) under Grant Number JP25ama121001 (Support Number 4257). This work was also supported by the Japan Society for the Promotion of Science (JSPS) Grant-in-Aid for Scientific Research (A) 24H00505 to T.N. and N.T.), Japan Science and Technology Agency (JST) PRESTO (JPMJPR21EA o N.T.), JST FOREST (JPMJFR2310 to N.T.), JST SPRING, Japan (JPMJSP2106 and JPMJSP2180 to H.L.), and Japan Society for the Promotion of Science Research Fellowships for Young Scientists 24KJ1042 to K.T.).

H.L. gratefully acknowledges scholarship support from the Amano Institute of Technology during 2023– 2026.

We thank the Materials Analysis Division, Core Facility Center, Research Infrastructure Management Center, Institute of Science Tokyo, for access to and technical support with TEM instrumentation.

OpenAI tools were used to assist with English-language editing. The authors have reviewed the manuscript and take full responsibility for its content.

## Supplementary information

**Figure S1.**
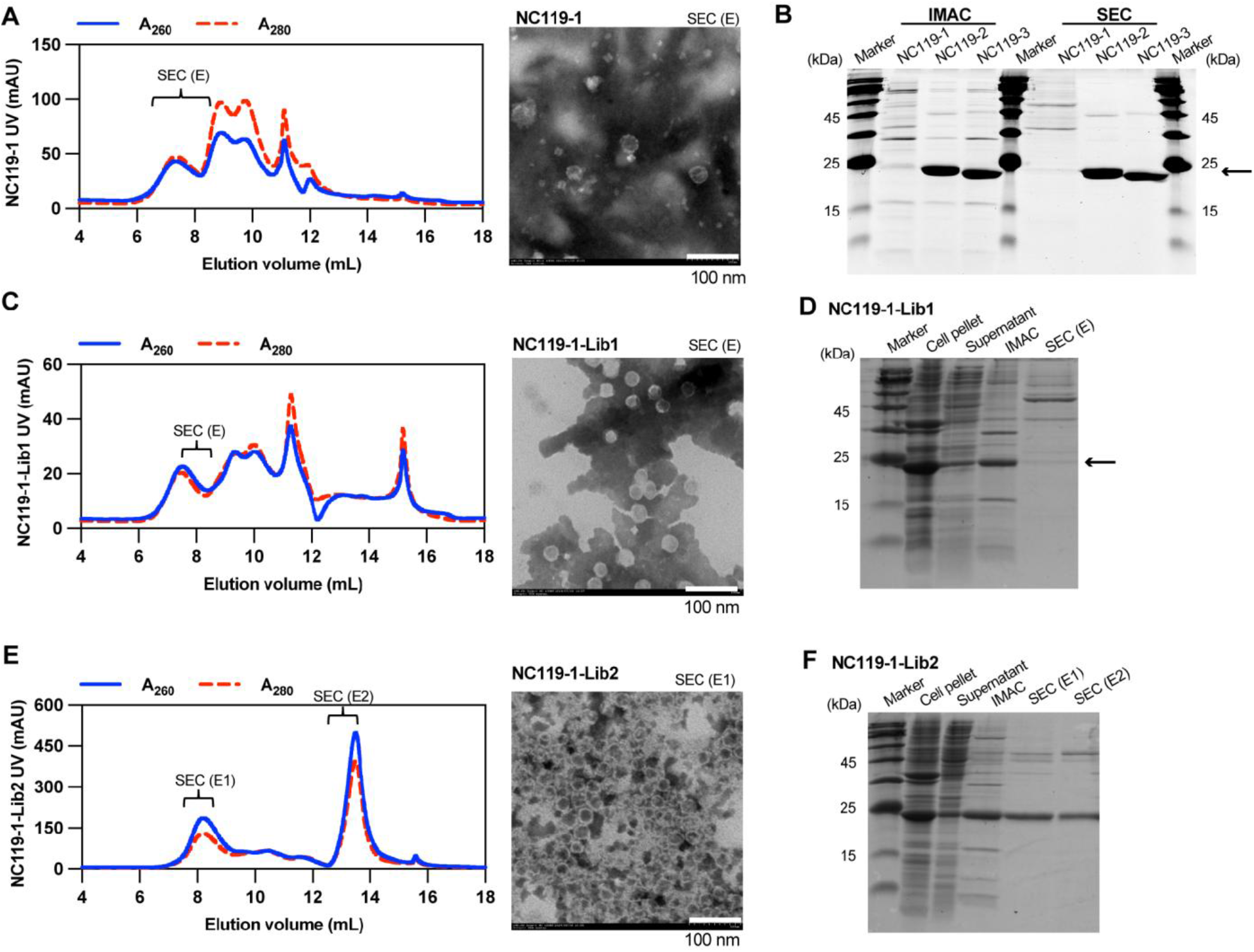
Initial characterization of NC119-1 and laboratory evolution. **(A)** Left: size-exclusion chromatography (SEC) profile of NC119-1. Solid blue and dashed red lines indicate absorbance at 260 nm (A260) and 280 nm (A280), respectively. Right: negative-stain transmission electron microscopy (nsTEM) image of SEC purified NC119-1. **(B)** SDS-PAGE analysis of NC119-1, NC119-2, and NC119-3 after immobilized metal affinity chromatography (IMAC) and SEC purification. The arrow indicates the expected position of the nucleocapsid protein monomer (∼21 kDa). **(C)** Left: SEC profile of NC119-1-Lib1. Right: nsTEM image of SEC purified NC119-1-Lib1. **(D)** SDS-PAGE analysis of the initial error-prone PCR library, NC119-1-Lib1, following expression in *E. coli*, IMAC purification and SEC purification. The arrow indicates the expected position of the nucleocapsid protein. The IMAC-purified sample was used for subsequent Benzonase treatment and RNA extraction. **(E)** Left: SEC profile of NC119-1-Lib2. Right: nsTEM images of NC119-1-Lib2 SEC eluates E1 (7.5–8.5 mL)**. (F)** SDS-PAGE analysis of the Benzonase-selected library NC119-1-Lib2 during purification. Cell pellet, supernatant, IMAC eluate, and eluates E1 (7.5–8.5 mL) and E2 (12.5–13.5 mL) are shown. Scale bars, 100 nm.

**Figure S2.**
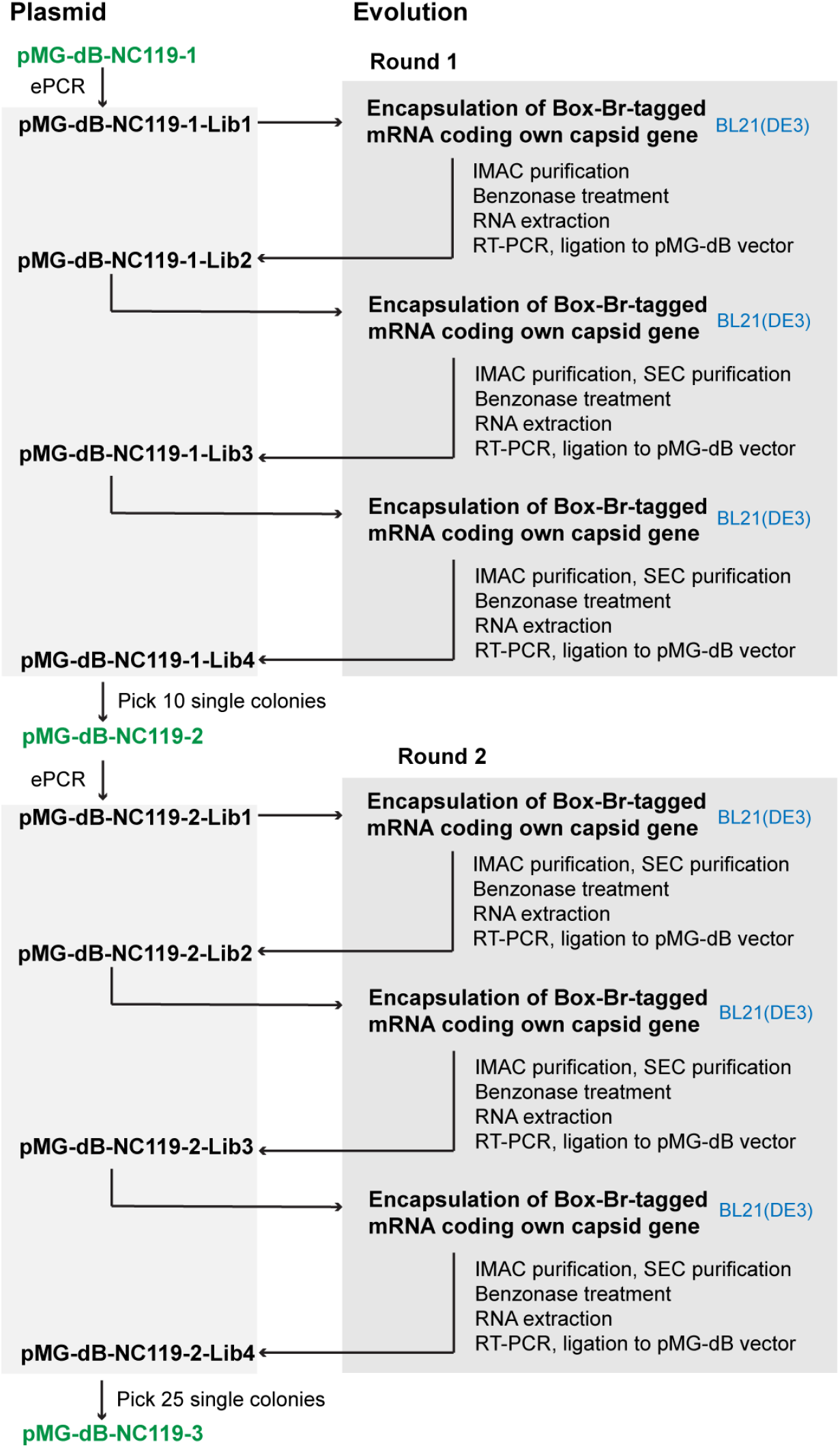
Overview of the directed evolution of NC119 nucleocapsids. Experimental details are described in the Experimental Section of the main text.

**Figure S3.**
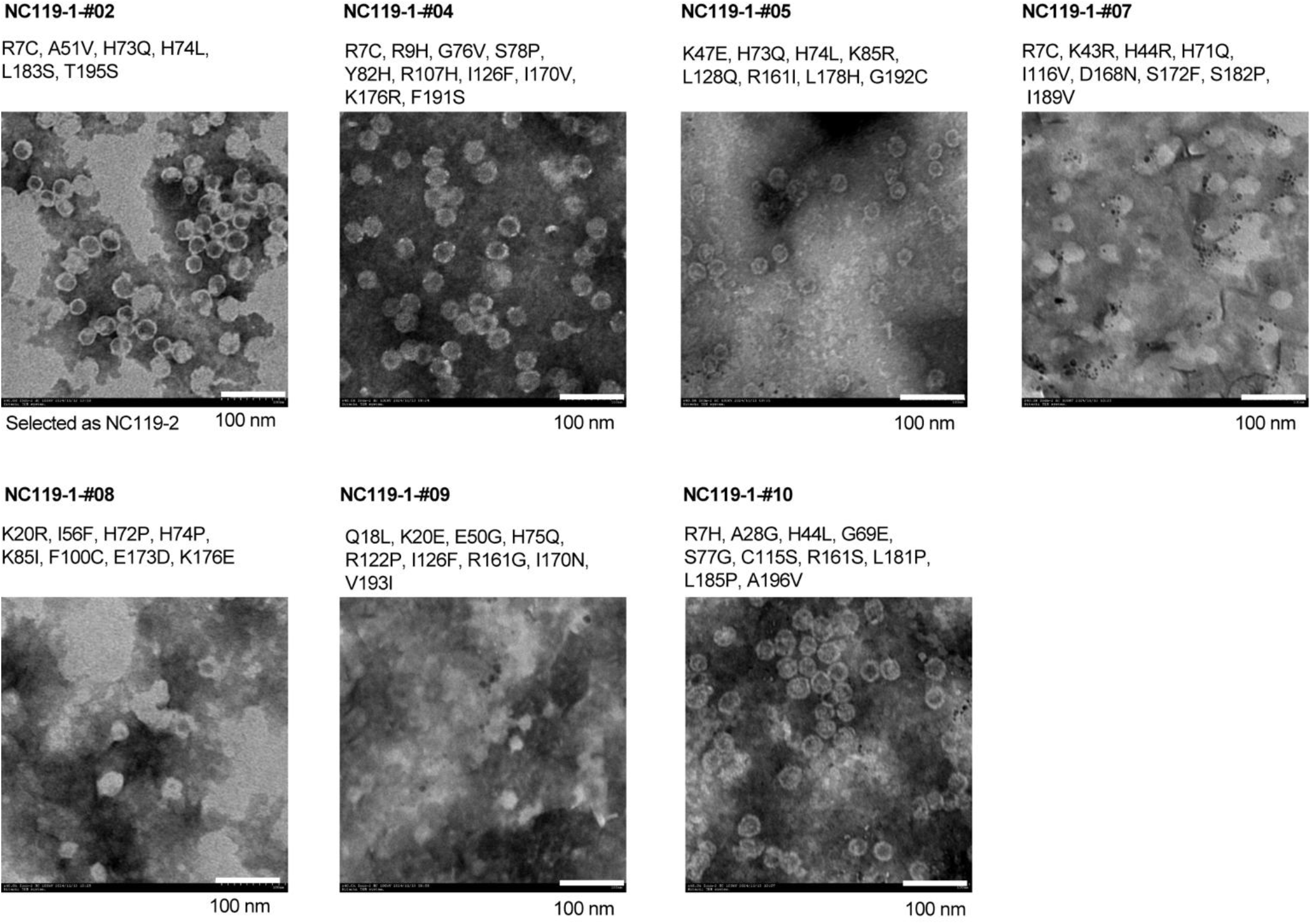
Assembly screening of individual variants after the first round of NC119 evolution. Ten individual colonies were randomly selected after three cycles of Benzonase treatment and analyzed by sequencing. Detailed nucleotide substitutions of the isolated variants are provided in Table S1. Seven variants with unique nucleotide sequences were identified, and their amino-acid substitutions relative to NC119-1 are shown together with representative nsTEM images. NC119-1#02 was selected as NC119-2 for the subsequent round of evolution. Scale bars, 100 nm.

**Figure S4.**
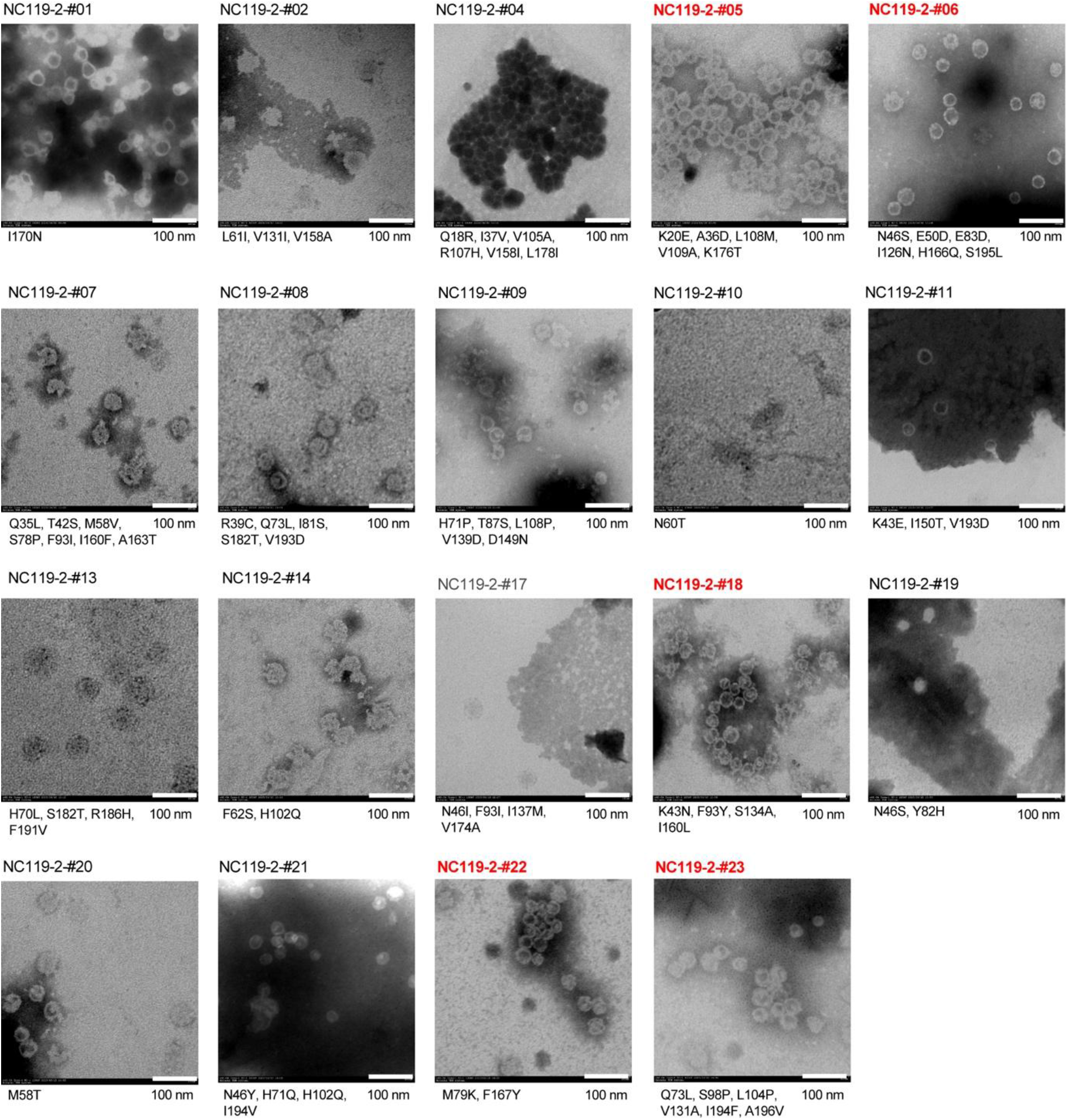
nsTEM analysis of individual variants after the second round of NC119 evolution. The 19 variants containing missense mutations were individually expressed in *E. coli*, purified by IMAC, and screened by nsTEM. Variants selected for further characterization in Figure S5 are highlighted in red. Scale bars, 100 nm.

**Figure S5.**
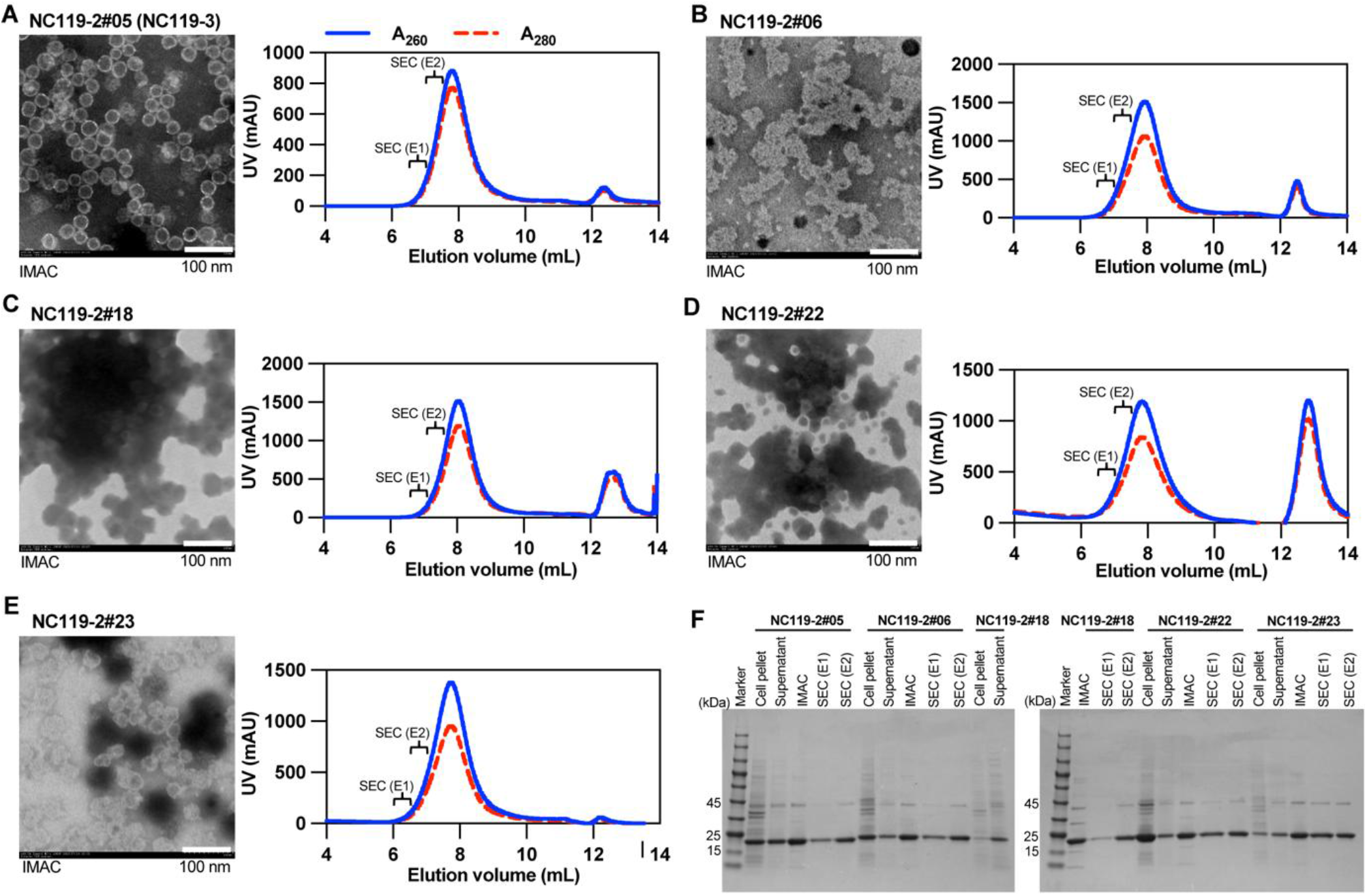
Further characterization of selected variants from the second round of NC119 evolution. **(A–E)** nsTEM images after IMAC purification and SEC profiles of five selected variants. The samples shown here were prepared in an independent expression experiment from those in Figure S4. Solid blue and dashed red lines indicate A260 and A280, respectively. SEC fractions E1 and E2 are indicated. Fractions eluted early were collected with the aim of recovering larger cages, although cages of different sizes were not clearly resolved by SEC. **(F)** SDS-PAGE analysis of the five variants during expression and purification. The NC protein migrated at approximately 21 kDa. Scale bars, 100 nm.

**Figure S6.**
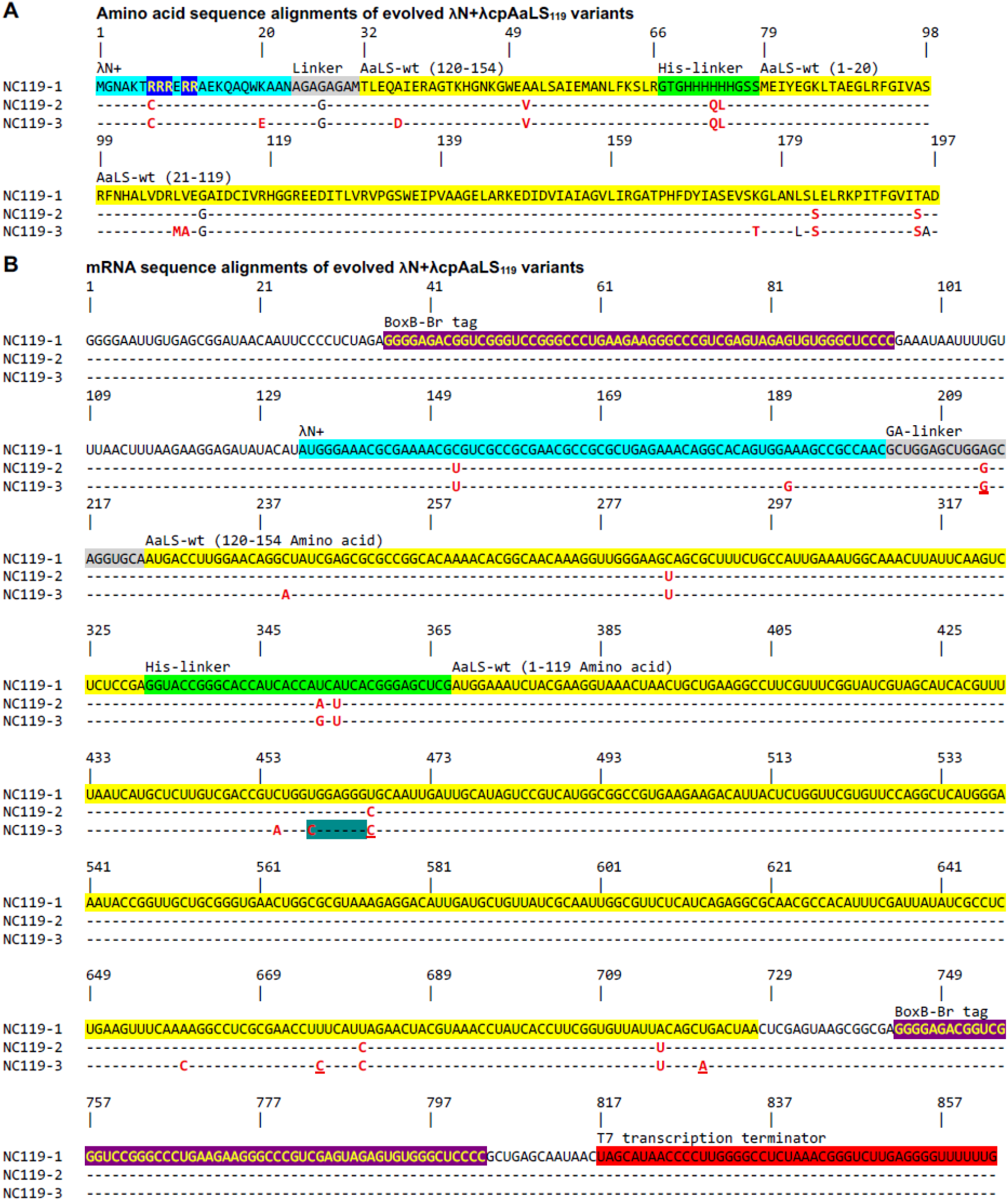
Amino acid and mRNA sequence alignments of evolved λN+cpAaLS119 variants. **(A)** Conserved residues are indicated by dashes, amino acid substitutions by red letters, and unchanged residues encoded by codons containing synonymous mutations by black letters. Amino acid positions are numbered according to the full-length λN+cpAaLS119 sequence, consistent with the numbering in Figure 2A. **(B)** Comparison of the variants with the λN+cpAaLS119 (NC119-1) mRNA sequence containing flanking BoxB-Br tags (top). Conserved nucleotides are indicated by dashes, and mutated nucleotides by red letters. Red underlining indicates synonymous mutations that do not alter the encoded amino acid. Blue highlighting marks the site where mutations eliminate a BoxB-like sequence motif (URxRxRR, where R denotes A or G and x denotes any nucleotide).

**Figure S7.**
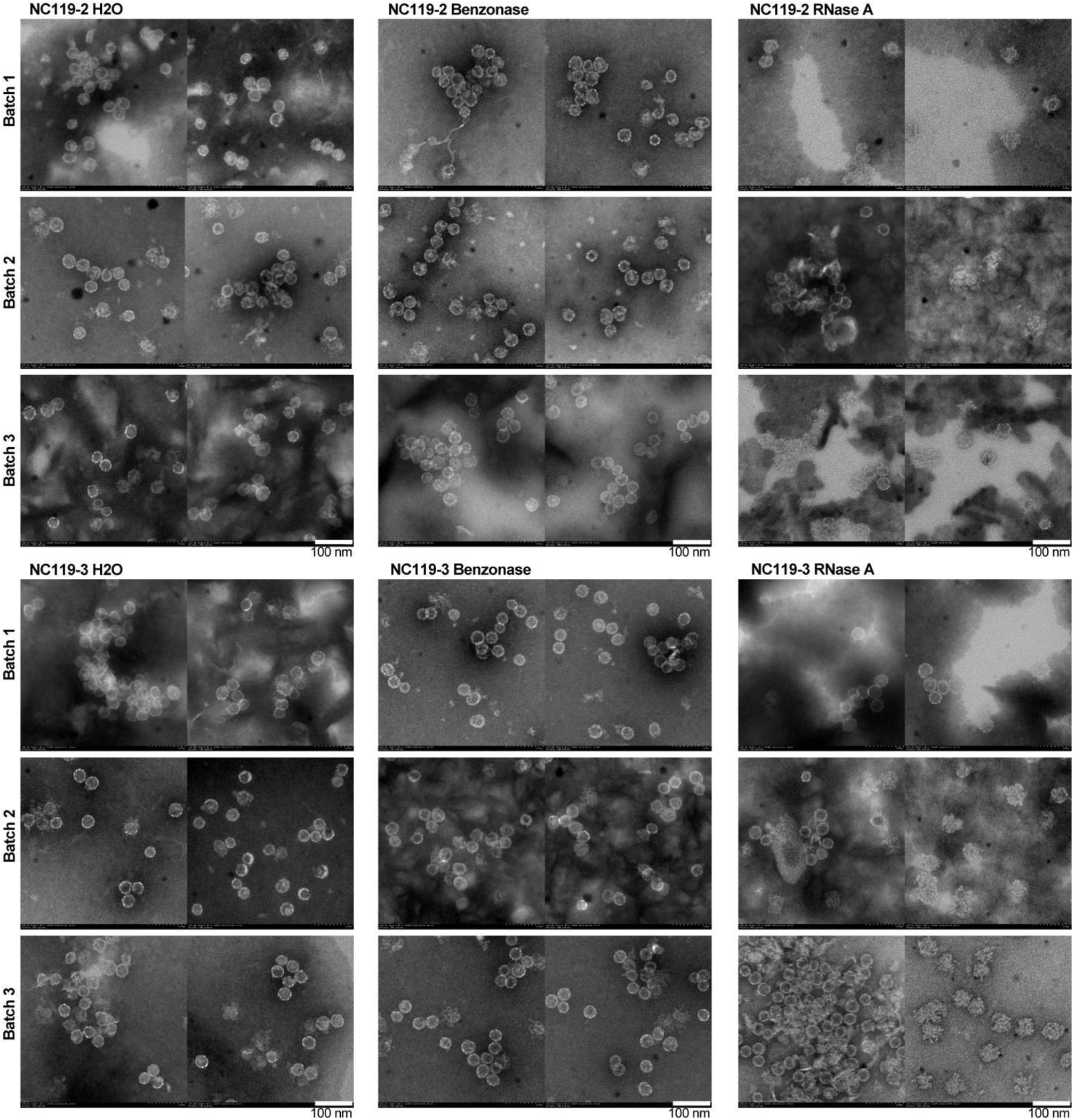
Morphology of evolved nucleocapsids following nuclease challenge. Negative-stain TEM (nsTEM) images of NC119-2 and NC119-3 following a 1 h incubation at 37°C with water (control), Benzonase, or RNase A. Two representative images are shown for each condition from each of three independent batches. Scale bars, 100 nm.

**Figure S8.**
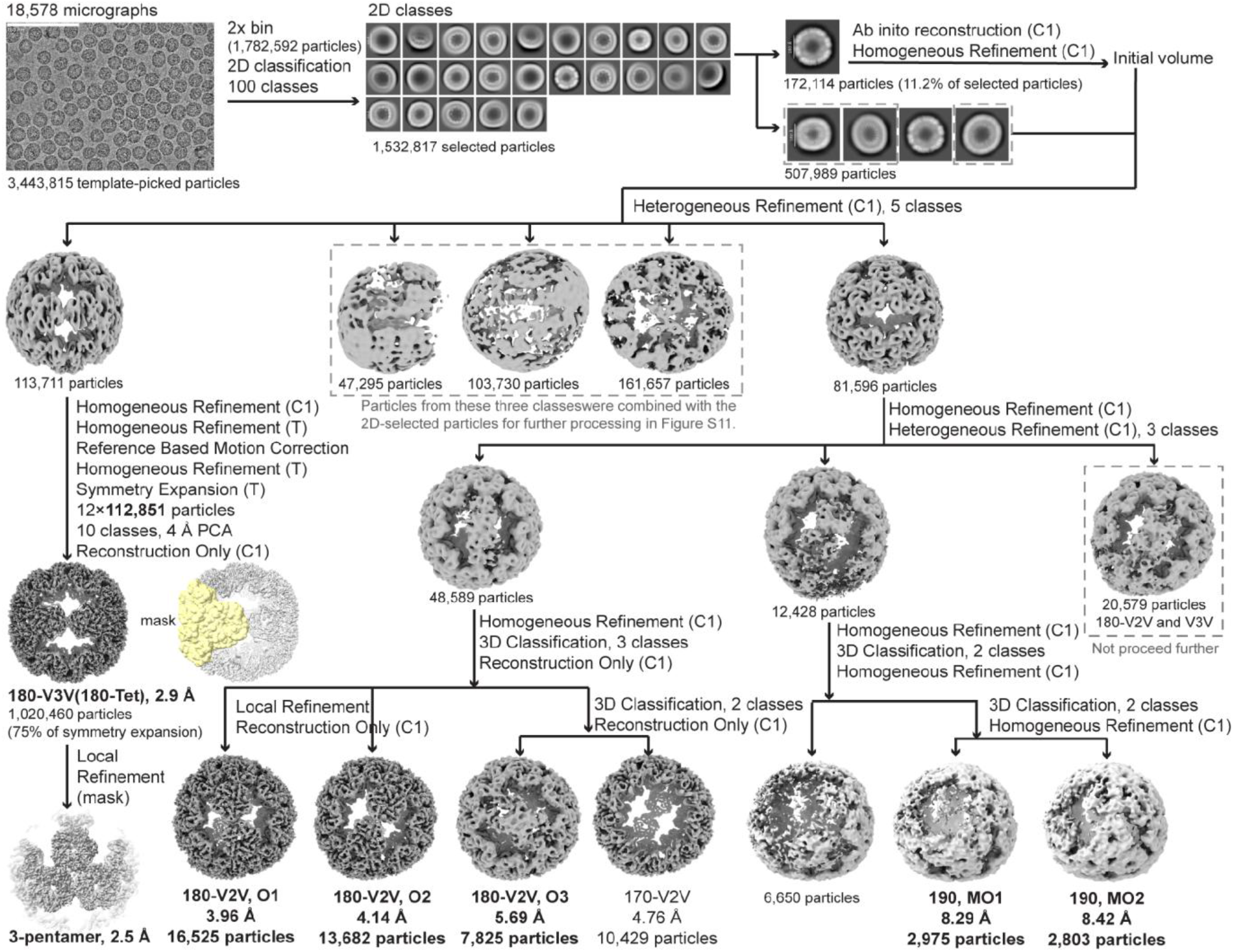
Cryo-EM processing workflow for the NC119-3 180-subunit family and minor enlarged branches. An initial three-dimensional cage reconstruction obtained from a preliminary set of picked particles was used to generate two-dimensional projection templates with the Create Templates job in cryoSPARC. Using these templates, particle picking from 18,578 micrographs yielded 3,443,815 particles. After twofold-binned extraction, 1,782,592 particles were classified into 100 two-dimensional classes, and 1,532,817 selected particles entered structural analysis. A 172,114-particle subset (11.2% of selected particles) generated an initial C1 volume. This volume was then used as a reference for heterogeneous refinement of a broader set of 507,989 particles to reduce the risk of excluding alternative architectures. Tetrahedral refinement, symmetry expansion, and focused classification yielded the 2.90 Å 180-V3V reconstruction, termed 180-Tet because the overall cage exhibits tetrahedral symmetry. Local refinement using a mask encompassing a three-pentamer module yielded a 2.50 Å reconstruction of this region. The 180-V2V branch yielded O1, O2, and O3 reconstructions with the particle counts and resolutions shown, together with a 170-V2V reconstruction that was not analyzed further or deposited in the EMDB. A minor lower-resolution branch produced two approximately 190-subunit reconstructions, designated 190 MO1 and 190 MO2.

**Figure S9.**
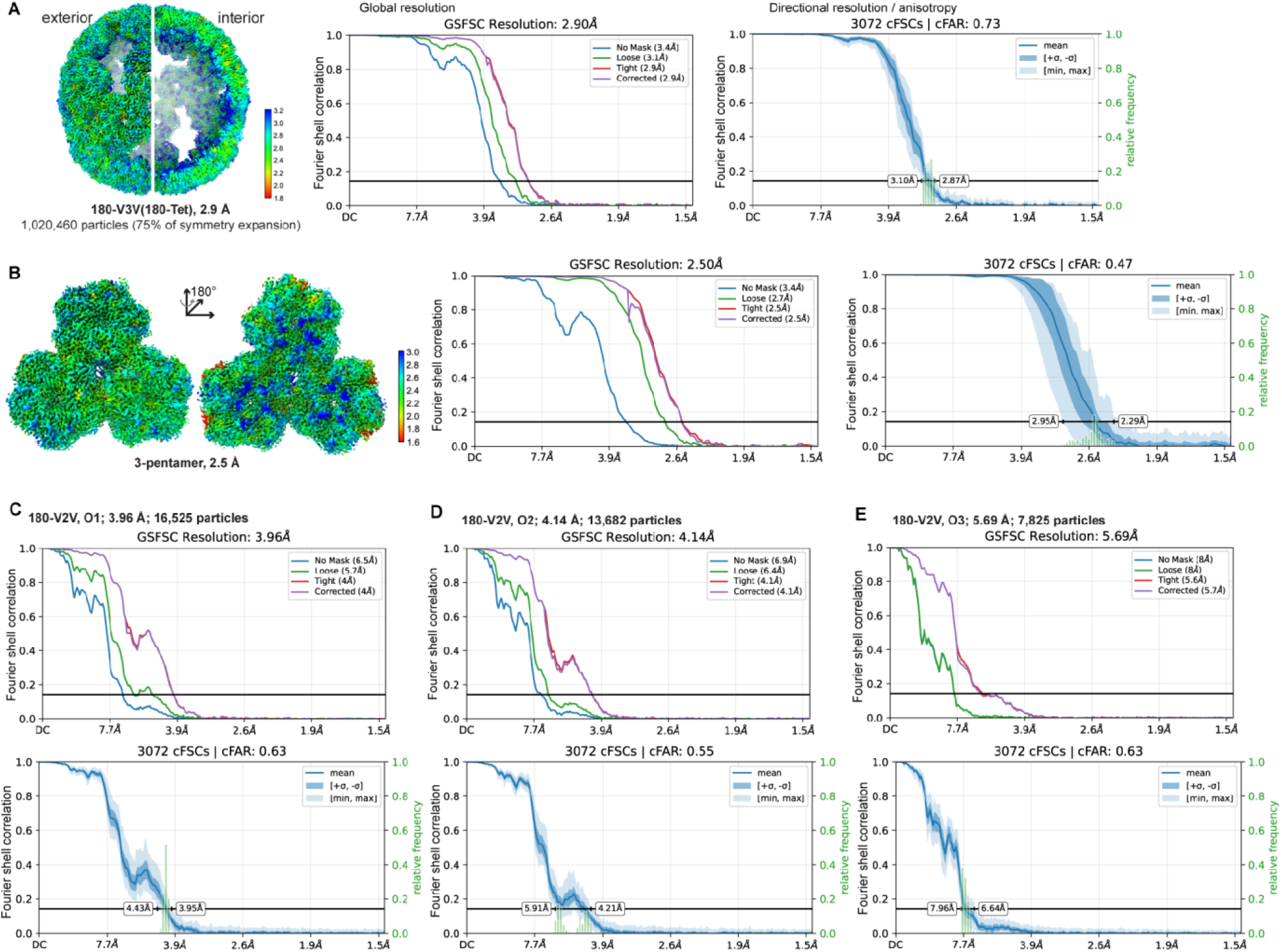
Resolution assessment of NC119-3 180-mer cryo-EM reconstructions. **(A)** Local-resolution map of the 180-V3V (180-Tet) reconstruction, shown in exterior and cutaway views, together with the gold-standard Fourier shell correlation (GS-FSC) curve and directional conical FSC (cFSC) analysis. The 180-V3V reconstruction reached a global resolution of 2.90 Å. **(B)** Local-resolution maps of the locally refined three-pentamer module shown in two views related by a 180° rotation, together with the corresponding GS-FSC and directional cFSC analyses. Local refinement yielded a resolution of 2.50 Å. **(C–E)** GS-FSC and directional cFSC analyses of the 180-V2V O1, O2, and O3 reconstructions, with global resolutions of 3.96, 4.14, and 5.69 Å, respectively. Resolution values were determined using the FSC = 0.143 criterion.

**Figure S10.**
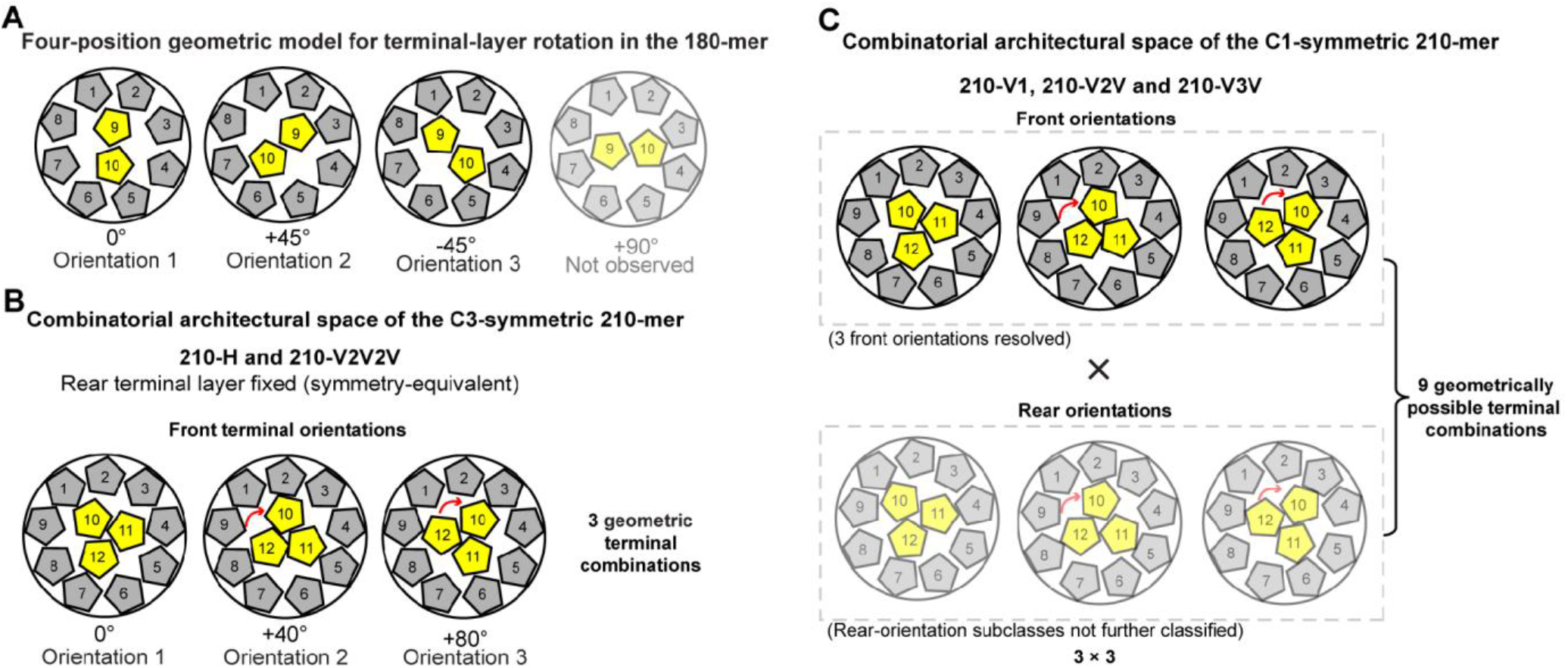
Geometric models of terminal-module orientations in the 180- and 210-subunit architectures. **(A)** Four-position geometric model of central pentamer-pair orientations in the 180-mer. With the rear terminal layer aligned to a common reference, the two central pentamers of the front terminal layer adopt three resolved orientations, corresponding approximately to 0° (O1), +45° (O2), and −45° (O3) relative to O1. A fourth orientation at approximately +90° is geometrically possible but was not observed. **(B)** Geometric model of the C3-symmetric 210-H and 210-V2V2V architectures. With the rear terminal layer fixed in a symmetry-equivalent orientation, the front terminal module can adopt three orientations, corresponding approximately to 0°, +40°, and +80°. **(C)** Combinatorial architectural space of the C1-symmetric 210-V1, 210-V2V, and 210-V3V architectures. Three orientations of the front terminal module were resolved, while the rear terminal module remained heterogeneous and its orientation subclasses were not further classified. The three possible front orientations combined with three geometrically possible rear orientations define nine geometrically possible terminal-orientation combinations. Angles shown are schematic geometric relationships rather than measured rigid-body rotation angles.

**Figure S11.**
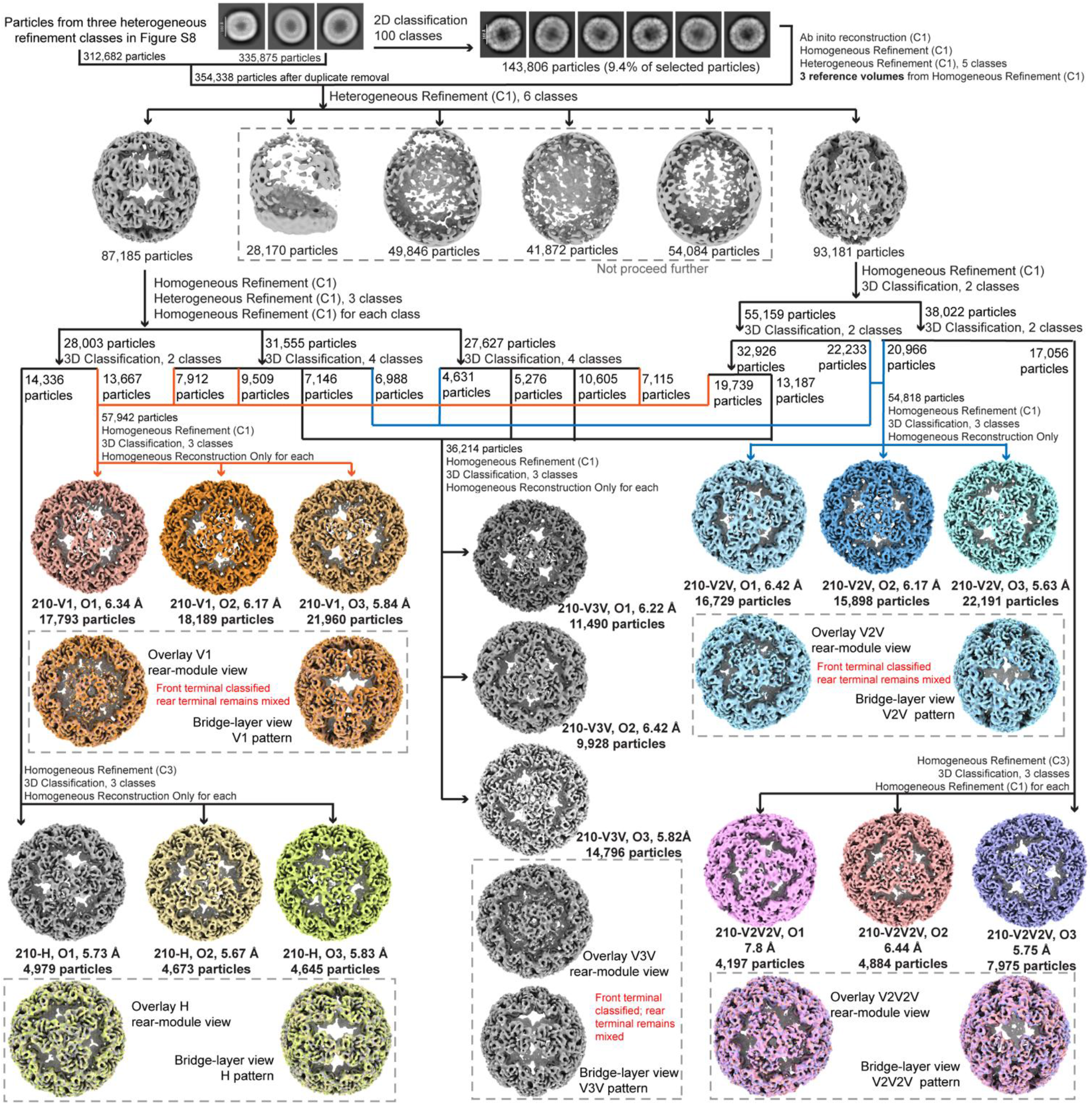
Cryo-EM classification workflow for the NC119-3 210-subunit family. Three 2D classes containing 335,875 particles were further classified, yielding 143,806 particles for ab-initio reconstruction and C1 refinement. Subsequent classification generated three reference volumes for heterogeneous refinement. The original 335,875 particles were combined with 312,682 particles from the three 3D classes boxed in Figure S8. After duplicate removal, 354,338 particles underwent heterogeneous refinement into six classes. Further refinement and 3D classification identified five middle-layer bridge patterns. Final maps show O1–O3 orientations of 210-V1, 210-H, 210-V2V, 210-V3V, and 210-V2V2V, with resolutions and particle counts indicated. Dashed boxes highlight rear-terminal modules and middle-layer bridge patterns. For 210-V1, 210-V2V, and 210-V3V, O1–O3 distinguish front-terminal orientations, while rear-terminal orientations remain mixed; the nine possible terminal-orientation combinations were not individually resolved. C1 and C3 indicate the symmetry settings used.

**Figure S12.**
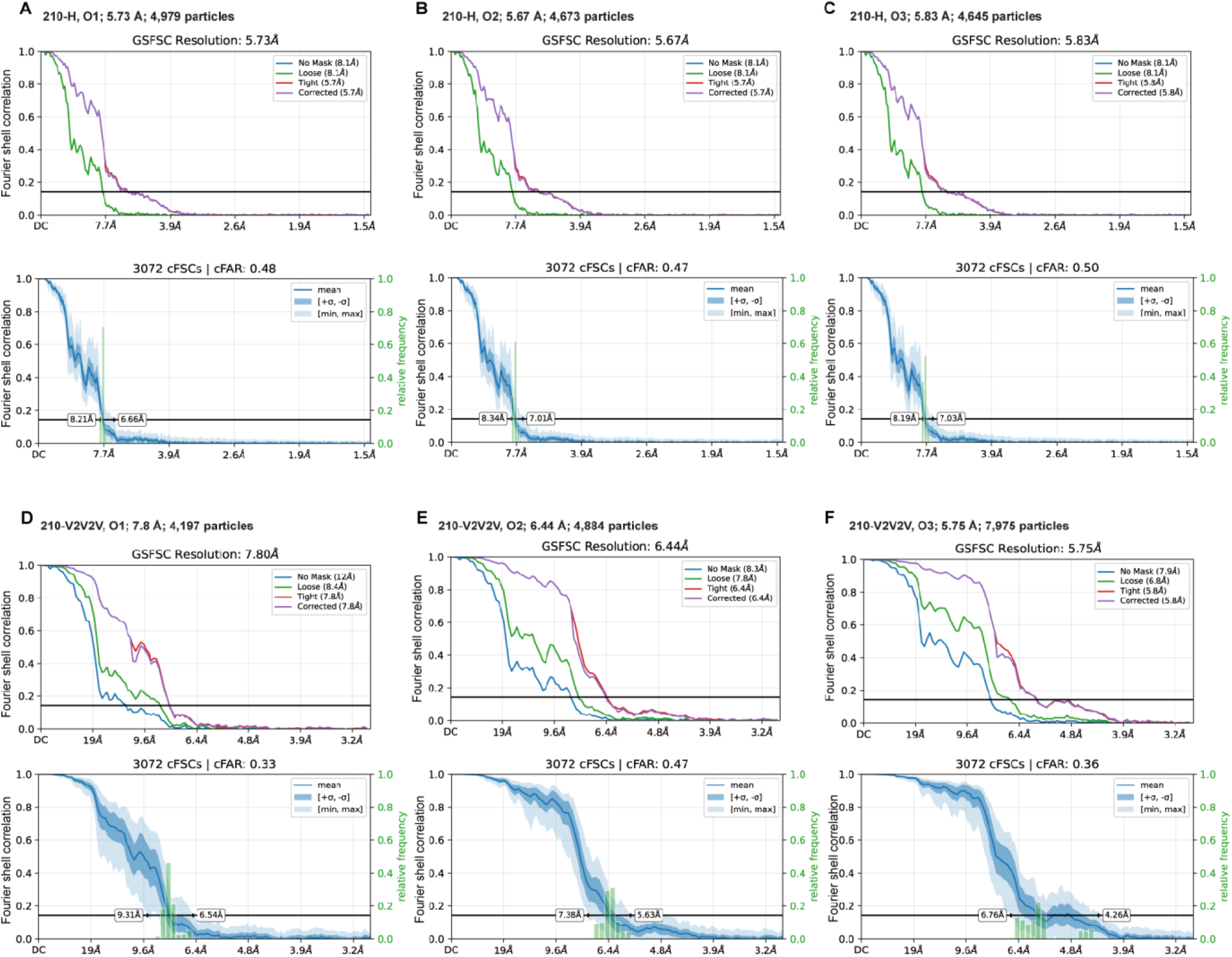
Resolution assessment of C3-symmetric 210-mer cryo-EM reconstructions. **(A–C)** Gold-standard Fourier shell correlation (GS-FSC) and directional conical FSC (cFSC) analyses of 210-H O1, O2, and O3, respectively. **(D–F)** Corresponding analyses of 210-V2V2V O1, O2, and O3. Particle numbers and global resolutions are indicated in each panel. Global resolutions were determined using the FSC = 0.143 criterion.

**Figure S13.**
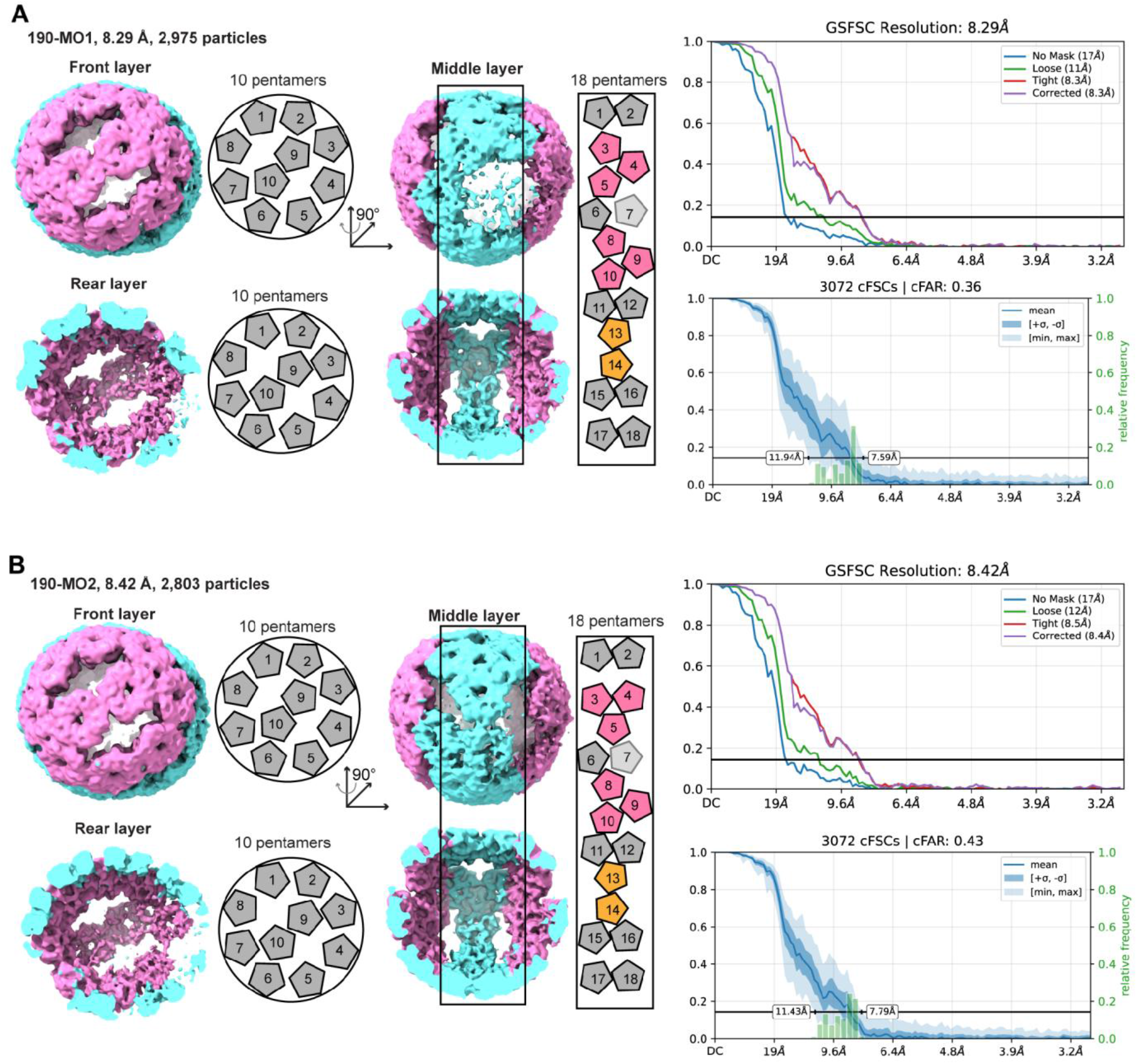
Structural organization and resolution assessment of the minor approximately 190-subunit architecture. **(A)** Cryo-EM reconstruction of 190-MO1 at 8.29 Å from 2,975 particles. The cage is interpreted as comprising 38 pentamers, with 10 pentamers in each terminal layer and 18 pentamers in the middle layer. Whole-cage and rear-terminal-layer views are shown alongside schematic representations of the terminal and middle layers. The middle-layer schematics indicate proposed three-pentamer bridges alongside paired-pentamer bridges. The corresponding gold-standard Fourier shell correlation (GS-FSC) and directional conical FSC (cFSC) analyses are shown on the right. **(B)** Cryo-EM reconstruction of 190-MO2 at 8.42 Å from 2,803 particles, displayed and analyzed as in (A). The proposed pentamer arrangements represent approximate interpretations of the maps, given the limited resolution and particle numbers. Global resolutions were determined using the FSC = 0.143 criterion.

**Figure S14.**
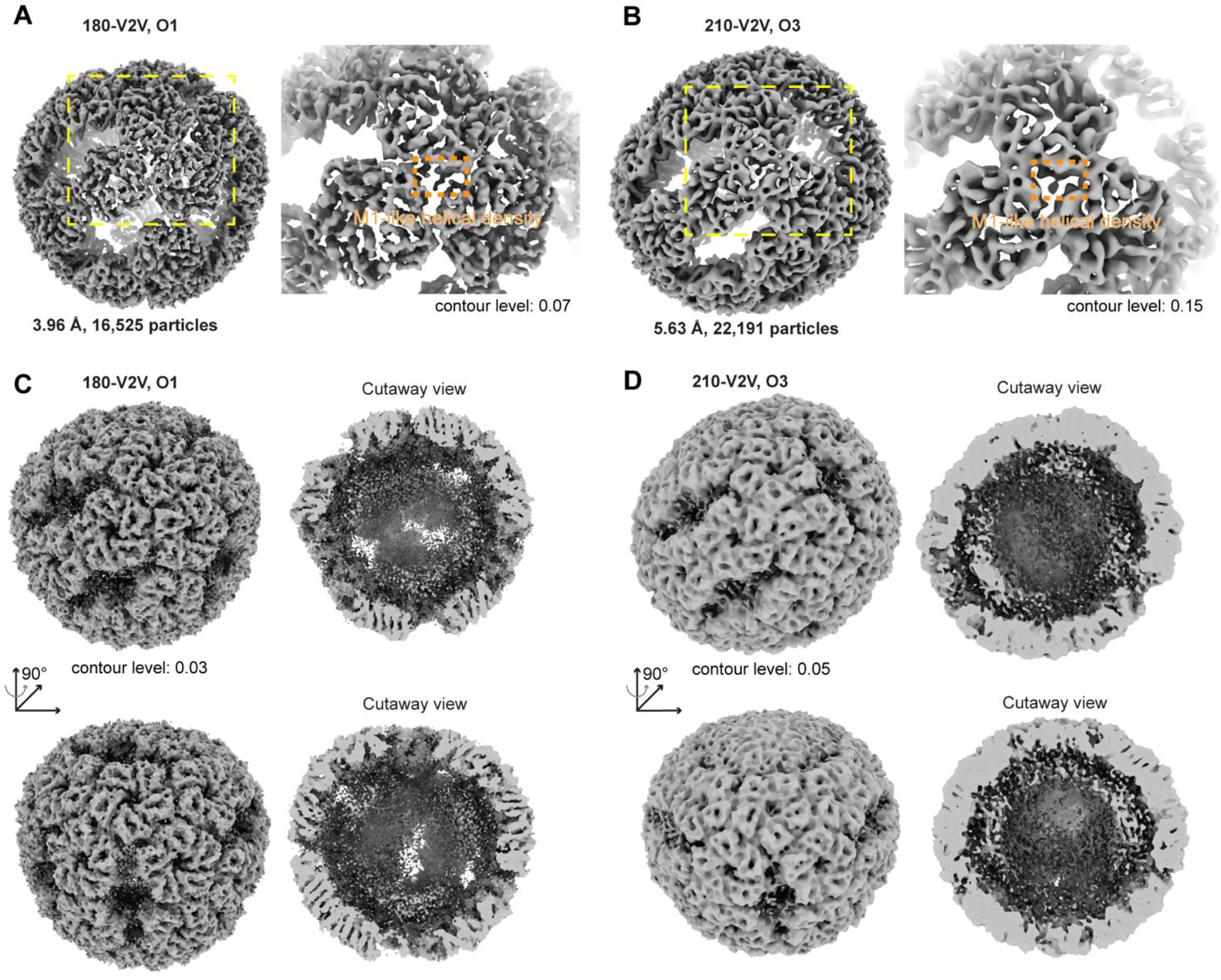
M1-like helical features and diffuse internal density in representative C1 reconstructions of NC119-3 cages. **(A, B)** Whole-cage views and enlarged regions of the 180-V2V O1 reconstruction (3.96 Å; 16,525 particles) and the 210-V2V O3 reconstruction (5.63 Å; 22,191 particles), respectively. Yellow dashed boxes indicate the enlarged regions, and orange dashed boxes highlight extended helical densities resembling the N-terminal helix of the M1 protomer at three-pentamer junctions. Maps are displayed at contour levels of 0.07 (A) and 0.15 (B). **(C, D)** Whole-cage and corresponding cutaway views of the same reconstructions at lower contour levels of 0.03 (C) and 0.05 (D), revealing diffuse density within the cage interiors. Two views related by a 90° rotation are shown for each reconstruction. The internal density may correspond in part to encapsulated RNA, although contributions from unresolved protein segments cannot be excluded. Both reconstructions were calculated without imposed symmetry (C1). The handedness of the 210-V2V O3 map was inverted in UCSF ChimeraX to match that of the 180-mer atomic model.

**Figure S15.**
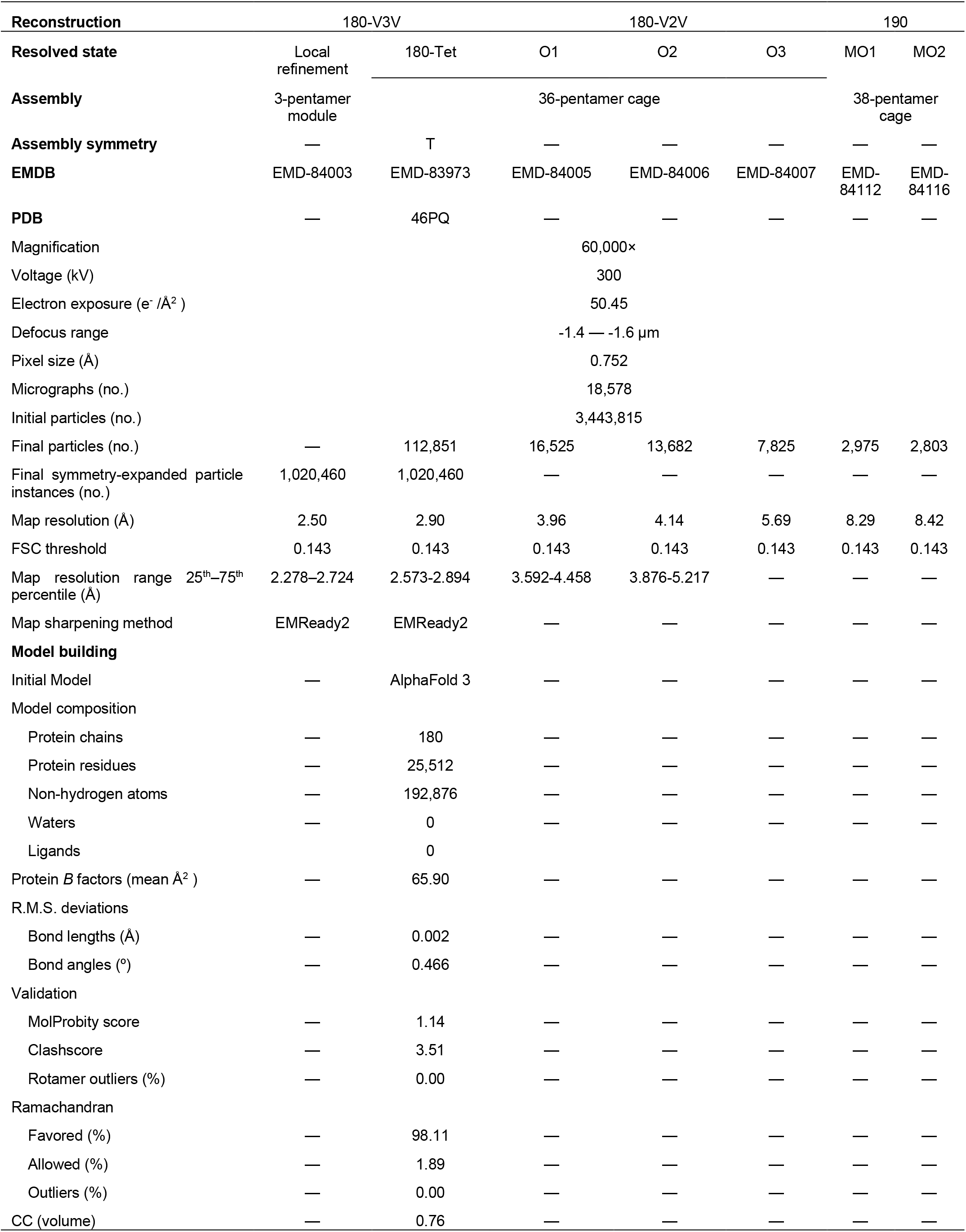
Cryo-EM reconstruction and validation statistics for the 180- and ∼190-subunit NC119-3 cages. Statistics include the locally refined three-pentamer module, 180-V3V (180-Tet), three 180-V2V orientations, and two tentative ∼190-subunit reconstructions. Model statistics refer to the full-cage 180-V3V structure. Both original and EMReady2-processed cryoSPARC maps were used for model building. Final Phenix refinement used the EMReady2-processed map, whereas validation used the original map. For 180-V3V, particle instances represent symmetry-expanded orientations retained after focused classification, not independent particles. Map resolutions were determined at FSC = 0.143. Dashes indicate unavailable or inapplicable data.

**Figure S16.**
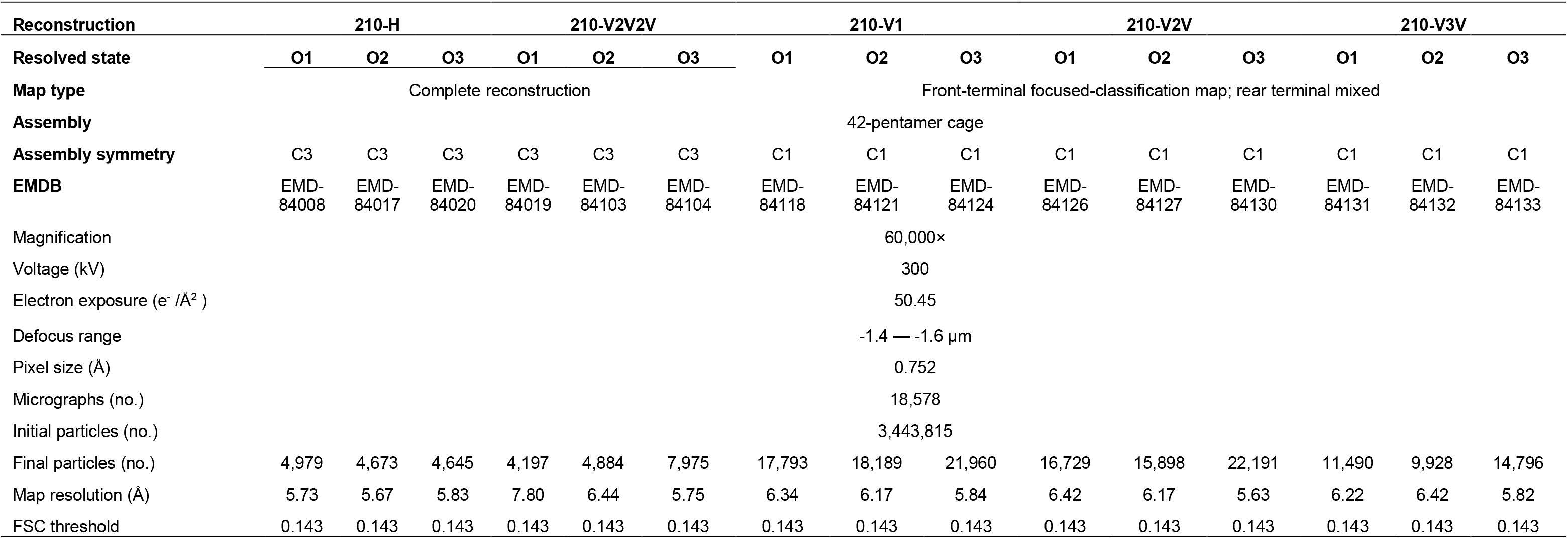
Cryo-EM data collection and reconstruction statistics for the 210-subunit family. Statistics are provided for the O1–O3 orientation classes of the 210-H, 210-V2V2V, 210-V1, 210-V2V, and 210-V3V architectures. The 210-H and 210-V2V2V reconstructions were refined with C3 symmetry. For 210-V1, 210-V2V, and 210-V3V, focused classification resolved the front-terminal orientations, while the rear-terminal orientations remained mixed. These nine classes were subsequently reconstructed in C1 without further alignment. Map resolutions were determined using the FSC = 0.143 criterion.

## References

1) Aumiller, W. M., Jr.; Uchida, M.; Douglas, T. Protein cage assembly across multiple length scales. Chem. Soc. Rev. 2018, 47 (10), 3433–3469. DOI: 10.1039/c7cs00818j.

2) Edwardson, T. G. W.; Levasseur, M. D.; Tetter, S.; Steinauer, A.; Hori, M.; Hilvert, D. Protein cages: From fundamentals to advanced applications. Chem. Rev. 2022, 122 (9), 9145–9197. DOI: 10.1021/acs.chemrev.1c00877.

3) Yeates, T. O. Geometric principles for designing highly symmetric self-assembling protein nanomaterials. Annu. Rev. Biophys. 2017, 46, 23–42. DOI: 10.1146/annurev-biophys-070816-033928.

4) Sharma, M.; Biela, A. P.; Kowalczyk, A.; Borzęcka-Solarz, K.; Piette, B. M. A. G.; Gaweł, S.; Bishop, J.; Kukura, P.; Benesch, J. L. P.; Imamura, M.;, et al. Shape-morphing of an artificial protein cage with unusual geometry induced by a single amino acid change. ACS Nanosci. Au 2022, 2 (5), 404–413. DOI: 10.1021/acsnanoscienceau.2c00019.

5) Sasaki, E.; Böhringer, D.; van de Waterbeemd, M.; Leibundgut, M.; Zschoche, R.; Heck, A. J.; Ban, N.; Hilvert, D. Structure and assembly of scalable porous protein cages. Nat. Commun. 2017, 8, 14663. DOI: 10.1038/ncomms14663.

6) Jorda, J.; Leibly, D. J.; Thompson, M. C.; Yeates, T. O. Structure of a novel 13 nm dodecahedral nanocage assembled from a redesigned bacterial microcompartment shell protein. Chem. Commun. 2016, 52 (28), 5041–5044. DOI: 10.1039/c6cc00851h.

7) Liang, S.; Butaney, K.; de Castro Assumpção, D.; Jung, J.; Kennedy, N. W.; Tullman-Ercek, D. Synthetic rewiring of virus-like particles via circular permutation enables modular peptide display and protein encapsulation. ACS Nano 2025, 19 (45), 39168–39180. DOI: 10.1021/acsnano.5c12293.

8) Osuna, J.; Pérez-Blancas, A.; Soberón, X. Improving a circularly permuted TEM-1 β-lactamase by directed evolution. Protein Eng. 2002, 15 (6), 463–470. DOI: 10.1093/protein/15.6.463.

9) Fang, J.; Chen, Z.; Liu, Y.; Zhang, W.-B. Directed evolution of a protein catenane reveals how noncanonical topology modulates sequence–property relationships. J. Am. Chem. Soc. 2026, 148 (40), 43280–43288. DOI: 10.1021/jacs.6c11777.

10) Zhang, X.; Meining, W.; Fischer, M.; Bacher, A.; Ladenstein, R. X-ray structure analysis and crystallographic refinement of lumazine synthase from the hyperthermophile Aquifex aeolicus at 1.6 Å esolution: Determinants of thermostability revealed from structural comparisons. J. Mol. Biol. 2001, 306 (5), 1099–1114. DOI: 10.1006/jmbi.2000.4435.

11) Wörsdörfer, B.; Woycechowsky, K. J.; Hilvert, D. Directed evolution of a protein container. Science 2011, 331 (6017), 589–592. DOI: 10.1126/science.1199081.

12) Azuma, Y.; Herger, M.; Hilvert, D. Diversification of protein cage structure using circularly permuted subunits. J. Am. Chem. Soc. 2018, 140 (2), 558–561. DOI: 10.1021/jacs.7b10513.

13) Terasaka, N.; Azuma, Y.; Hilvert, D. Laboratory evolution of virus-like nucleocapsids from nonviral protein cages. Proc. Natl. Acad. Sci. U. S. A. 2018, 115 (21), 5432–5437. DOI: 10.1073/pnas.1800527115.

14) Caspar, D. L.; Klug, A. Physical principles in the construction of regular viruses. Cold Spring Harbor Symp. Quant. Biol. 1962, 27, 1–24. DOI: 10.1101/sqb.1962.027.001.005.

15) Tetter, S.; Terasaka, N.; Steinauer, A.; Bingham, R. J.; Clark, S.; Scott, A. J. P.; Patel, N.; Leibundgut, M.; Wroblewski, E.; Ban, N.;, et al. Evolution of a virus-like architecture and packaging mechanism in a repurposed bacterial protein. Science 2021, 372 (6547), 1220–1224. DOI: 10.1126/science.abg2822.

16) Koziej, L.; Fatehi, F.; Aleksejczuk, M.; Byrne, M. J.; Heddle, J. G.; Twarock, R.; Azuma, Y. Dynamic assembly of pentamer-based protein nanotubes. ACS Nano 2025, 19 (9), 8786–8798. DOI: 10.1021/acsnano.4c16192.

17) Porterfield, J. Z.; Zlotnick, A. A simple and general method for determining the protein and nucleic acid content of viruses by UV absorbance. Virology 2010, 407 (2), 281–288. DOI: 10.1016/j.virol.2010.08.015.

18) Filonov, G. S.; Kam, C. W.; Song, W.; Jaffrey, S. R. In-gel imaging of RNA processing using Broccoli reveals optimal aptamer expression strategies. Chem. Biol. 2015, 22 (5), 649–660. DOI: 10.1016/j.chembiol.2015.04.018.

19) Beren, C.; Dreesens, L. L.; Liu, K. N.; Knobler, C. M.; Gelbart, W. M. The effect of RNA secondary structure on the self-assembly of viral capsids. Biophys. J. 2017, 113 (2), 339–347. DOI: 10.1016/j.bpj.2017.06.038.

20) Cadena-Nava, R. D.; Comas-Garcia, M.; Garmann, R. F.; Rao, A. L.; Knobler, C. M.; Gelbart, W. M. Self-assembly of viral capsid protein and RNA molecules of different sizes: Requirement for a specific high protein/RNA mass ratio. J. Virol. 2012, 86 (6), 3318–3326. DOI: 10.1128/jvi.06566-11.

21) Tajima, K.; Sakai, Y.; Terasaka, N. Cargo-directed assembly of nonviral nucleocapsid with controlled size. ACS Synth. Biol. 2026, 15 (3), 1178–1186. DOI: 10.1021/acssynbio.5c00881.

22) Seitz, I.; Saarinen, S.; Kumpula, E.-P.; McNeale, D.; Anaya-Plaza, E.; Lampinen, V.; Hytönen, V. P.; Sainsbury, F.; Cornelissen, J. J. L. M.; Linko, V.;, et al. DNA-origami-directed virus capsid polymorphism. Nat. Nanotechnol. 2023, 18 (10), 1205–1212. DOI: 10.1038/s41565-023-01443-x.

23) Valegârd, K.; Murray, J. B.; Stonehouse, N. J.; van den Worm, S.; Stockley, P. G.; Liljas, L. The three-dimensional structures of two complexes between recombinant MS2 capsids and RNA operator fragments reveal sequence-specific protein-RNA interactions. J. Mol. Biol. 1997, 270 (5), 724–738. DOI: 10.1006/jmbi.1997.1144.

24) Dykeman, E. C.; Stockley, P. G.; Twarock, R. Dynamic allostery controls coat protein conformer switching during MS2 phage assembly. J. Mol. Biol. 2010, 395 (5), 916–923. DOI: 10.1016/j.jmb.2009.11.016.

25) Rolfsson, Ó.; Toropova, K.; Ranson, N. A.; Stockley, P. G. Mutually-induced conformational switching of RNA and coat protein underpins efficient assembly of a viral capsid. J. Mol. Biol. 2010, 401 2), 309–322. DOI: 10.1016/j.jmb.2010.05.058.

26) Beck, T.; Tetter, S.; Künzle, M.; Hilvert, D. Construction of matryoshka-type structures from supercharged protein nanocages. Angew. Chem., Int. Ed. 2015, 54 (3), 937–940. DOI: 10.1002/anie.201408677.

27) Schindelin, J.; Arganda-Carreras, I.; Frise, E.; Kaynig, V.; Longair, M.; Pietzsch, T.; Preibisch, S.; Rueden, C.; Saalfeld, S.; Schmid, B.;, et al. Fiji: An open-source platform for biological-image analysis. Nat. Methods 2012, 9 (7), 676–682. DOI: 10.1038/nmeth.2019.

28) Yonekura, K.; Maki-Yonekura, S.; Naitow, H.; Hamaguchi, T.; Takaba, K. Machine learning-based real-time object locator/evaluator for cryo-EM data collection. Commun. Biol. 2021, 4 (1), 1044. DOI: 10.1038/s42003-021-02577-1.

29) Mastronarde, D. N. Advanced data acquisition from electron microscopes with SerialEM. Microsc. Microanal. 2018, 24 (S1), 864–865. DOI: 10.1017/S1431927618004816.

30) Punjani, A.; Rubinstein, J. L.; Fleet, D. J.; Brubaker, M. A. cryoSPARC: Algorithms for rapid unsupervised cryo-EM structure determination. Nat. Methods 2017, 14 (3), 290–296. DOI: 10.1038/nmeth.4169.

31) Abramson, J.; Adler, J.; Dunger, J.; Evans, R.; Green, T.; Pritzel, A.; Ronneberger, O.; Willmore, L.; Ballard, A. J.; Bambrick, J.;, et al. Accurate structure prediction of biomolecular interactions with AlphaFold 3. Nature 2024, 630 (8016), 493–500. DOI: 10.1038/s41586-024-07487-w.

32) Cao, H.; Zhu, Y.; Li, T.; Chen, J.; He, J.; Wang, X.; Huang, S.-Y. EMReady2: Improvement of cryo-EM and cryo-ET maps by local quality-aware deep learning with Mamba. Nat. Commun. 2026, 17 (1), 6493. DOI: 10.1038/s41467-026-71794-1.

33) Croll, T. I. ISOLDE: A physically realistic environment for model building into low-resolution electron-density maps. Acta Crystallogr., Sect. D: Struct. Biol. 2018, 74, 519–530. DOI: 10.1107/s2059798318002425.

34) Pettersen, E. F.; Goddard, T. D.; Huang, C. C.; Meng, E. C.; Couch, G. S.; Croll, T. I.; Morris, J. H.; Ferrin, T. E. UCSF ChimeraX: Structure visualization for researchers, educators, and developers. Protein Sci. 2021, 30 (1), 70–82. DOI: 10.1002/pro.3943.

35) Emsley, P.; Lohkamp, B.; Scott, W. G.; Cowtan, K. Features and development of Coot. Acta Crystallogr., Sect. D: Biol. Crystallogr. 2010, 66, 486–501. DOI: 10.1107/s0907444910007493.

36) Liebschner, D.; Afonine, P. V.; Baker, M. L.; Bunkóczi, G.; Chen, V. B.; Croll, T. I.; Hintze, B.; Hung, L. W.; Jain, S.; McCoy, A. J.;, et al. Macromolecular structure determination using X-rays, neutrons and electrons: Recent developments in Phenix. Acta Crystallogr., Sect. D: Struct. Biol. 2019, 75, 861–877. DOI: 10.1107/s2059798319011471.

